# A causal role for estradiol in human reinforcement learning

**DOI:** 10.1101/2020.02.18.954982

**Authors:** Sebastijan Veselic, Gerhard Jocham, Christian Gausterer, Bernhard Wagner, Miriam Ernhoefer-Reßler, Rupert Lanzenberger, Claus Lamm, Christoph Eisenegger, Annabel Losecaat Vermeer

## Abstract

The sex hormone estrogen is hypothesized to play a key role in human cognition via its interactions with the dopaminergic system. Work in rodents has shown that estrogen’s most potent form, estradiol, impacts striatal dopamine functioning predominately via increased D1-receptor signalling, while human work has suggested that high estradiol levels are associated with altered reward sensitivity. Here, we addressed two fundamental questions: 1) whether estradiol causally alters reward sensitivity in men, and 2) whether this effect of estradiol is moderated by individual variation in polymorphisms of dopaminergic genes. To test this, we performed a double-blind placebo-controlled administration study in which hundred men received either a single dose of estradiol (2 mg) or placebo. We found that estradiol administration increased reward sensitivity, which was moderated by baseline dopamine. This was observed in choice behaviour and increased learning rates. These results confirm a causal role of estradiol in reinforcement learning in men that is moderated by the striatal dopaminergic pathway.

## Introduction

Learning which action to select based on whether the outcome of that action is rewarded or not is a fundamental capacity required for adaptive behaviour. One neuromodulator that has long been linked to this capacity, known as reinforcement learning (RL), is dopamine ^1^. More recently, an additional biological substrate that has been suggested to influence RL via dopaminergic mechanisms is estrogen ^2^.

Estrogens are a class of steroid hormones important for healthy development, with estradiol being the most prevalent and potent form ^3,4^. Estradiol has gained traction as a compound that may impact human reward processing by amplifying dopamine signalling via the D1 receptor ^2^. The evidence for this hypothesis comes from two lines of work. From human work, we know that fluctuations in circulating estradiol levels are correlated with differences in midbrain BOLD responses, a key area where dopamine is released ^5–7^. Furthermore, a recent administration study in women showed that increased salivary estradiol levels led to larger reward prediction errors in the nucleus accumbens ^8^. From rodent work, we know that manipulating estradiol levels affects the striatal dopamine system in various ways that are best characterised as a net increase in overall dopamine signalling predominantly via the D1 receptor ^9–14^.

If increased estradiol levels in humans influence reward processing by increasing reward sensitivity through amplified dopamine signalling via the D1 receptor, then it should be possible to relate its effects to that of dopamine in RL^1,15–17^. Within this field, dopamine agonists and antagonists are often administered to better understand the mechanistic role of dopamine in RL ^18–24^. Complementing it with pharmacogenetics, where the effect of genetic variation on the administered drug is considered, has refined our understanding of their relationship ^19,22,25–27^. That is, causally manipulating dopamine levels in humans affects performance in RL tasks^27^ with these effects depending on individual differences in baseline dopamine levels ^18^.

Two key polymorphisms that lead to variation in baseline dopamine levels by impacting dopamine synthesis capacity and transmission are the dopamine transporter (DAT1) and catechol-O-methyltransferase (COMT) gene ^19,24,25^. Variation in the VNTR polymorphism of DAT1 and the val^158^met polymorphism of COMT correlates with performance on working memory and RL in humans ^19,24,25,28,29^. Therefore, if estradiol influences reward processing through dopaminergic mechanisms, then variation in the DAT1 and COMT genotype should moderate the effect of estradiol. Furthermore, we would predict it should be moderated by traits related to human reward sensitivity such as the ones measured by the BIS/BAS questionnaire ^30–33^.

Despite abundant evidence from rodent research showing a clear relationship between estradiol and dopamine, evidence in humans have been less conclusive. Namely, it has been shown that high endogenous estradiol levels were associated with increased ^5,7^ as well as decreased ^34^ performance on a variety of cognitive tasks that may have recruited different neural mechanisms. Thus, it remains unclear whether administering estradiol influences reward processing as would be expected by amplified D1-receptor signalling. Furthermore, although previous work on humans provided important insights, it had certain shortcomings such as small sample sizes, correlational study designs, and a lack of accounting for baseline differences in dopamine; for exceptions see ^8,34,35 25,36^. All studies thus far have also focussed on female samples who have, on average, higher endogenous levels of estradiol compared to men ^37^. These aspects are important for being able to establish a more precise role of estradiol in human reward processing which, therefore, remains an open question (for review see ^2^).

The aim of the present pharmacogenetic study was to address these gaps and investigate whether estradiol administration increases reward sensitivity in men and whether this effect is moderated by baseline differences in dopamine. To test this, we used a probabilistic RL task (Fig. 1A) where subjects had to choose between two options on each trial in order to maximize their earnings. Moreover, we aimed at providing a more conclusive and precise account of a dopamine-dependent basis of action through excluding several other candidate explanations (i.e. gene polymorphisms) that could have given rise to the obtained results and have so far been unaddressed in the literature (see Supplementary Materials).

**Fig. 1.**
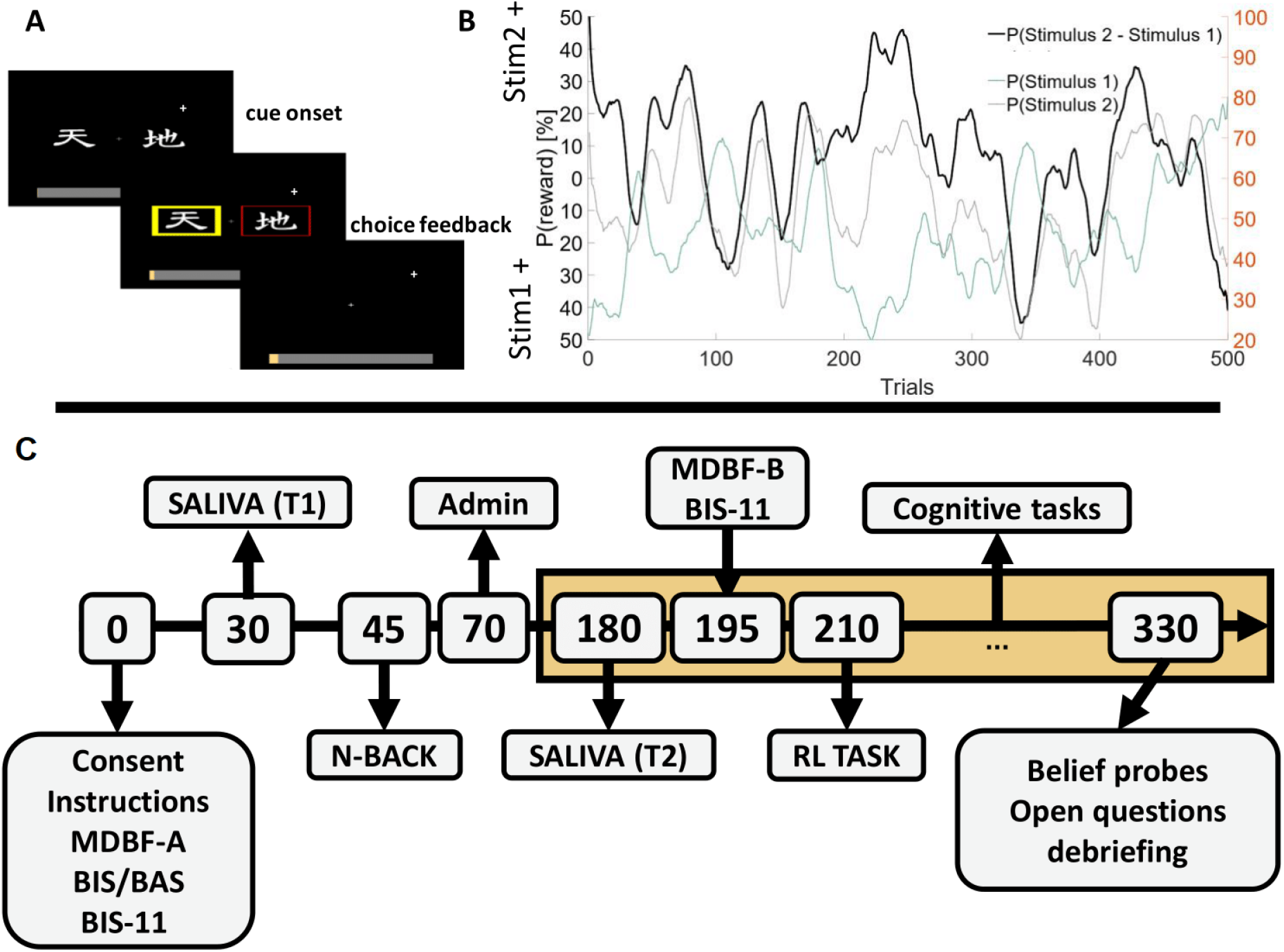
**A)** Outline of a trial of the RL task. Each trial started by the presentation of two options (henceforth option A and option B). Subjects were required to choose one of these options. After they made a choice, subjects were presented with feedback, with the chosen option indicated by a thicker frame and the not chosen option by a thinner frame. A yellow frame indicated the rewarded option, whereas a red frame indicated the unrewarded option. Importantly, both options A and B could yield a reward or no reward on the same trial. **B)** The probability of reward upon choice for each option (green and gray lines), which were determined by two independent random Gaussian walks, with the probability shown in percent on the right y-axis in orange. The black line shows the relative probability of reward for one option over the other, which corresponds to the difference in reward probability for option A and option B. On trials where the black line is reaching the top half of the y-axis, option A was more rewarding, and vice versa. **C)** The timeline of the test session. Values in brackets denote minutes from the onset of the test session. We first collected consent and questionnaire data, which was followed by a baseline saliva sample (T1) and the N-BACK task. After administration of estradiol or placebo, subjects were required to rest for two hours before we collected the second saliva sample (T2) and assessed subjects’ mood and impulsivity via questionnaires. The RL task began 120 minutes post-administration. This was followed by three other cognitive tasks that are not the focus of the current paper. At the end of the test session, we probed subjects’ beliefs about the drug, the experiment, and debriefed them.

We hypothesized that estradiol administration would influence reward processing by increasing reward sensitivity observed through subjects’ choice behaviour ^8^. We further predicted this would be revealed through computational modelling. Finally, we predicted that the behavioural and computational effects would be moderated by the DAT1 polymorphism, as observed in previous dopamine administration work ^19,26^.

## Methods and materials

### Subjects

We tested one hundred healthy young men between 19 and 34 years (*M*_*age*_ = 24.86, *SD* = 3.53) with a body mass index (BMI) between 19.3 and 31.5 (*M* = 24.45, *SD* = 2.86). Our screening procedure was based on previous work where pharmacokinetic data for a single 2 mg estradiol dose in topical form was obtained ^38^. Our sample size was in line with previous recommendations for the field ^2^. The study procedure was performed in accordance with the Declaration of Helsinki and approved by the Medical Ethics Committee of the University of Vienna (1918/2015).

### Measurement Instruments

#### Questionnaires

We assessed self-reported mood (German Multidimensial Mood State Questionnaire; MDBF^39^), individuals’ impulsiveness (Barratt Impulsiveness Scale; BIS-11^40^), and behavioural activation and inhibition (BIS/BAS^41^). Both BIS/BAS and BIS-11 scores have been previously found to correlate with reward learning ^42–45^ with the BIS/BAS specifically being related to reward sensitivity ^30,31^. In addition, we probed subjects’ beliefs about estradiol (e.g. whether they believed they received estradiol or a placebo, how certain they were about their belief, and whether they had noticed any subjective changes).

#### Hormone concentrations

We collected hormone samples via passive drool and stored them at −30 degrees Celsius. Saliva samples were analyzed for estrone and estradiol using gas chromatography tandem mass spectrometry (GC-MS/MS) and hydrocortisone including testosterone with liquid chromatography tandem mass spectrometry (LCMS/MS).

#### Genotyping

We collected DNA using sterile cotton buccal swabs (Sarstedt AG, Germany) and extracted it by applying the QIAamp DNA Mini kit (Qiagen, Germany). Repeat length polymorphisms (AR(CAG), AR(GGN), DAT1(VNTR), ERα(TA) and ERβ(CA)) were investigated by PCR with fluorescent-dye-labeled primers and capillary electrophoresis. The single base primer extension (SBE) method also known as minisequencing was applied for the typing of single nucleotide polymorphism (SNP) variants (Val158Met) in the COMT gene.

#### Experimental Tasks

##### Reinforcement Learning

We used a well-established probabilistic reinforcement learning task with two options ^17^. The task consisted of 500 trials, with a 10 second pause after the first 250 trials. There was no choice time-out. The two options had independently varying reward probabilities generated by Gaussian random walks (Fig. 1). Importantly, both options could be correct (i.e. rewarding - yellow frame) or incorrect (i.e. non-rewarding - reward frame) on any trial. Subjects received feedback for the chosen (thick frame) and unchosen (thin frame) option. Each correct choice was rewarded with 5 eurocents and added to their cumulative balance. Subjects also saw a bar fill up as they chose the correct option. Once the bar filled up, a 1 € coin was presented next to the bar indicating they had added 1 € to their cumulative balance.

##### Working memory capacity

We used an adapted version of the standard N-BACK task ^25^ with four conditions in total (0-BACK, 1-BACK, 2-BACK, 3-BACK). Each condition block had 20 trials which included 20% target, 65% nontarget, and 15% lure trials. Subjects were presented with a sequence of letters one-by-one. For each letter, they had to decide if the current letter was the same as the one presented *N* trials ago by pressing “R”, in case it was not the same they had to press “O”.

### Procedure

Subjects screened for BMI, a history of psychiatric disorders, concurrent involvement in other psychopharmacological studies, and chronic physical injuries, arrived to the lab on two separate days. On the first session, at 4.00 pm, we collected subjects’ responses on a battery of questionnaires, their DNA with a buccal swab for genotyping, and measured their BMI and body fat using a body composition monitor (Omron BF51).

On the second session (Fig. 1, timeline) after obtaining consent, subjects filled out the MDBF-A, BIS-11, and BIS/BAS. Twenty minutes after arrival, we obtained the first saliva sample (T1) to assess baseline hormone concentrations, followed by the N-BACK task. They were randomly assigned estradiol or placebo in a double-blind manner and self-administered a topical transparent gel containing either 2 mg of estradiol or a placebo. We waited two hours to allow estradiol levels to peak based on a previously established procedure ^38^. Fifteen minutes prior to the behavioural testing, subjects filled out MDBF-B, BIS-11, and provided a second saliva sample (T2).

The first behavioural task was the probabilistic reinforcement learning task, followed by three other tasks that were not the focus of this publication. After the behavioural testing, we probed subjects’ beliefs about the treatment and the tasks. At the end of the study, each subject was paid in accordance to their performance.

### Statistical analysis

#### Behavioural analysis

To determine the effect of estradiol on choice behaviour, we examined the cumulative difference in the options subjects chose and the percentage of trials on which there was a significant difference in the chosen option. As a measure of family-wise error control, we used permutation testing and Fisher-z-transformations.

We also computed choice accuracy because we predicted that if estradiol would influence reward processing by increasing reward sensitivity, this would result in increased accuracy and be moderated by differences in striatal dopamine functioning (i.e. through the DAT1 genotype) ^26^. Due to the same reasoning we computed how many trials subjects would pick the same option if they were rewarded for that option on trial *t* (staying). Accuracy and staying were statistically evaluated with general linear models.

In addition to the above metrics, we computed subjects’ choice autocorrelation. With this, we tested our hypothesis that increased reward prediction errors would, by definition, upregulate saliency of recent events ^8^. Behaviourally, this would mean subjects’ recent choices should influence what they will choose next more. We computed the relative contribution of choices made from *n* − 1 to *n* − 7 trials back (lags) on the current choice. We then compared the relative contribution of all previous choices (pure choice autocorrelation) and specifically choices when they were rewarded (choice autocorrelation as a function of reward). We then performed independent samples *t*-tests on individual lags that were z-scored within subject to assess statistical significance.

#### Computational modelling

To account for behaviour within a computational framework and relate our findings to the field exploring dopamine and its role in RL, we used computational modelling. We fit a series of Q-learning models with softmax choice rules. The winning model included a learning rate for positive and negative prediction errors, a temperature parameter, and an irreducible noise parameter (see Supplementary materials for model selection and parameter recovery). Our main hypothesis was that if estradiol would increase reward sensitivity, that should be captured by the learning rate, reflecting how strongly new information will be weighed and incorporated into the subjects’ subjective values.

## Results

Our sample was matched for age, height, visceral, and abdominal fat, BMI, working memory, self-reported impulsivity, behavioural inhibition and approach, and mood. As a manipulation check of our administration protocol, estradiol concentrations were significantly elevated in subjects who had received estradiol compared to placebo after (*W* = 1545, 95% CI [0.03, 1.87], *p* < .05), but not before administration (baseline: *W* = 1498, 95% CI [-0.05, 1.03], *p* = .09) and subjects’ beliefs about whether they had received estradiol or placebo did not correlate with the actual received drug (*r* = 0.02, *p* = .82; for further details see Supplementary Materials).

### Estradiol administration increases reward sensitivity

Our first hypothesis was that estradiol administration would increase reward sensitivity. Because reward size was kept constant, the only difference across trials was whether a choice was rewarded or not. Therefore, we predicted that increased reward sensitivity would be reflected in a systematic difference in the options subjects would choose across trials compared to placebo. To test this, we computed the average probability per group to select option A/B (i.e. expected chosen option) on each trial (Fig. 2A), subtracted these traces from each other, and plotted the cumulative choice difference across trials (Fig. 2B).

**Fig. 2.**
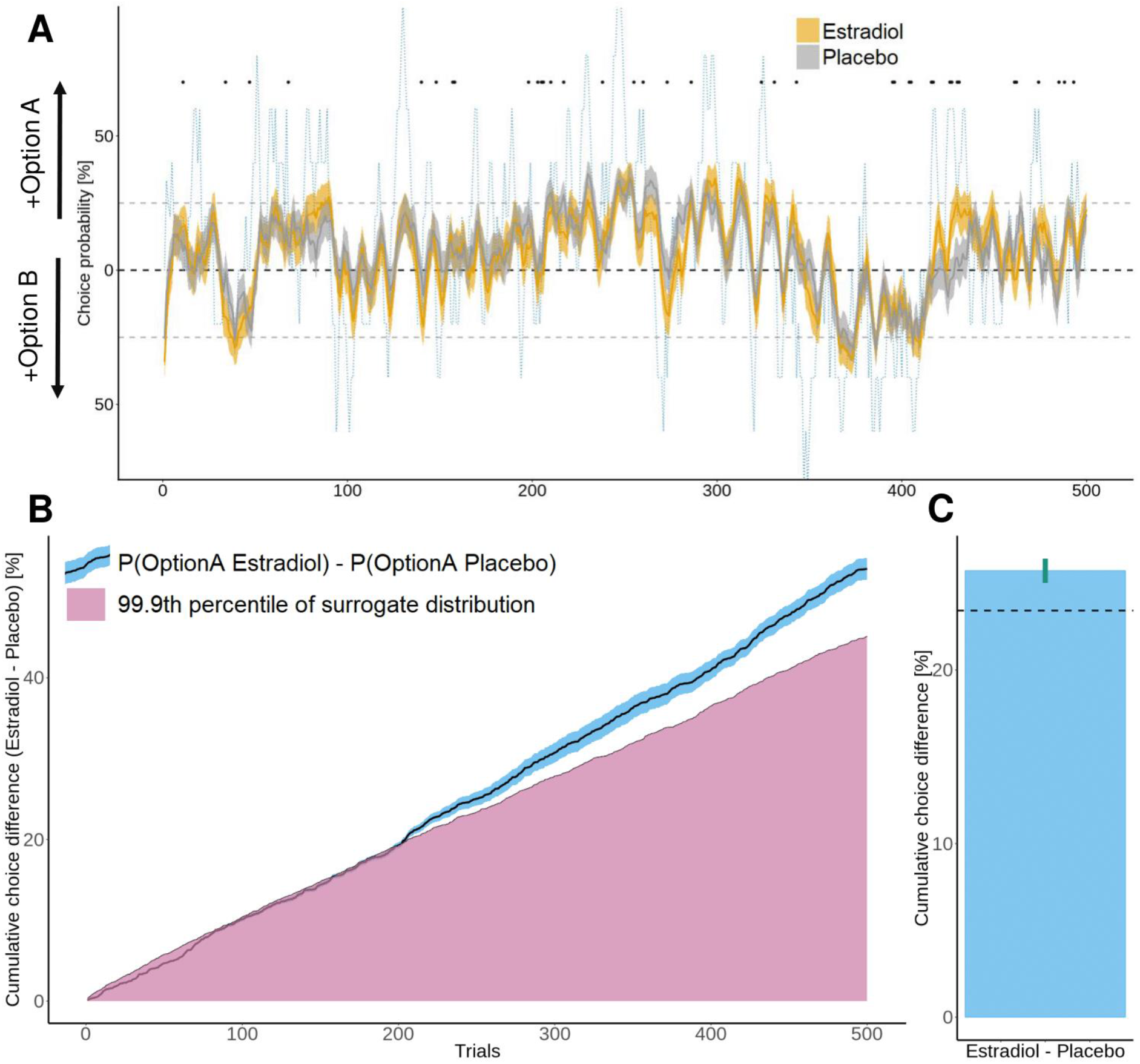
**A**) Relative choice probability for choosing option A (top of y-axis) vs. choosing option B (bottom of y-axis) for the estradiol (orange) and placebo (gray) group. Solid thick lines represent trial mean, shaded areas around the thick lines denote standard errors of the mean. The blue dotted line denotes the relative reward probability which was computed from the probability of option A (top of y-axis) minus probability of option B (bottom of y-axis). Horizontal gray dotted lines represent where subjects were on average 25% more likely to select option A (upper line) or option B (lower line). All time-series traces were smoothed with a 5-trial moving average for visual purposes. The black dots indicate trials where there was a statistically significant difference (*p* < .05) between the estradiol and placebo group. The number of significant trials was compared to a null distribution (see Methods and materials). **B**) Cumulative choice difference between the estradiol and placebo group over trials compared to the 99.9th^th^ percentile null distribution. The thick black line is the difference between the orange and gray lines presented in figure **A**, and the blue shaded area is the corresponding difference between the standard errors in **A**. The dark orange area denotes the space in which differences are not significant. Conversely, separation between the lines indicate statistical significance. **C)** Mean cumulative choice difference between the estradiol and placebo group collapsed across trials. The dashed line represents the mean cumulative choice difference of the 100^th^ percentile of the null distribution. Error bars indicate standard error of the mean.

The cumulative choice difference in the expected chosen option between both groups surpassed what would be expected by chance. This was tested by comparing the above results to the 99.9^th^ percentile of a null distribution (see Methods and materials) indicating what would be expected by chance (*M*_last trial_ = 53.48 %, *z*_last trial_ = 8.44, *p* < .001, threshold for 99.9^th^ percentile of the null: 46.20 %, Fig. 2B). The observed cumulative choice difference remained significant when collapsed across time (*M* = 25.72 ±0.69%, *z* = 5.80, *p* < .001, threshold for 99.9^th^ percentile of the null: *M* = 21.02 %, Fig. 2C). When we traced trials that contributed most to the cumulative difference, we observed a statistically significant difference in what both groups chose on 7.6 % of trials (black dots in Fig. 2A). In other words, estradiol administration caused subjects to choose a different option on 7.6 % of trials as compared to placebo (z = 5.37, *p* < .001, threshold for 99.9^th^ percentile of null distribution: 6.4 %).

However, estradiol administration did not only increase reward sensitivity in choice behaviour as described above. When subjects rated how strongly they observed the changing of reward probabilities of both options throughout the task on a scale from 1 to 100 during debriefing, those receiving estradiol explicitly reported that they observed reward probabilities across trials to change more strongly (*t*_(78.495)_ = 2.15, *95% CI* = [0.855, 22.61], *p* = 0.035, *d* = 0.48, *BF*_10_ = 3.28).

### Estradiol’s improvement in accuracy is moderated by DAT1 genotype and reward responsiveness

Following the observed systematic choice difference between both groups, we investigated whether this was reflected in group differences in accuracy. Both groups were equally accurate (*M*_*Estradiol*_ = 57.30 ±6.91, *M*_*Placebo*_ = 56.80 ±7.09, *t*_(97.94)_ = 0.36, *p* = .72, Fig. 3A), and responded equally fast (*M*_*Estradiol*_ = 0.61 sec ±0.11, *M*_*Placebo*_ = 0.62 *sec* ±0.09, *t*_(95.55)_ = 0.46, *p* = .65).

**Fig. 3.**
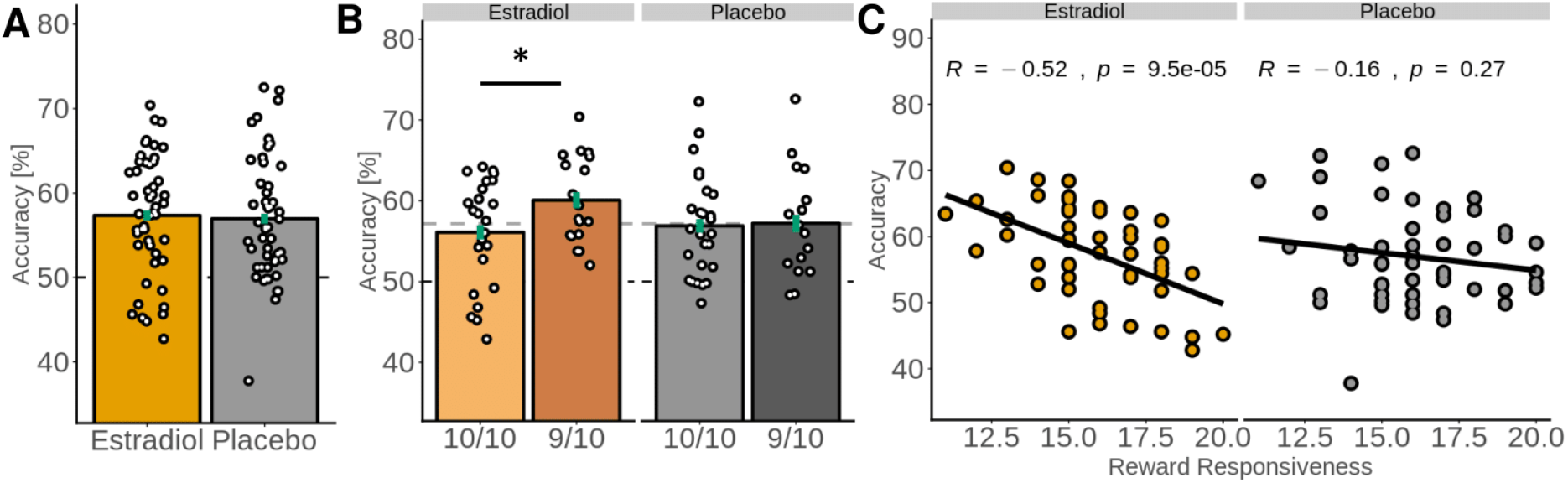
**A**) Mean accuracy split according to drug administration. **B**) Mean accuracy split according to drug administration and DAT1 polymorphism. **C**) Accuracy was moderated by reward responsiveness. Green error bars are standard errors of the mean. Dots represent individual subjects. The horizontal dotted line represents grand mean performance collapsed across groups to show the relative change for individual subgroups. * *p* < .05

However, we predicted that the effect of estradiol on accuracy will be moderated by individual differences in baseline striatal dopamine, as other work has shown interactive effects between task performance and dopamine-related genes ^18,46^. In our study, we used the DAT1 polymorphism as an index of striatal dopamine with the 9/10 and 10/10 genotypes being associated with high and low striatal dopamine, respectively ^47^.

A general linear model revealed an interaction between drug administration and DAT1 genotype on accuracy (*F*_(1, 69)_ = 4.10, *p* = .047, Ω^2^ = 0.037, Fig. 3B) while controlling for covariates (see Supplementary Materials). Pairwise comparisons revealed that estradiol administration increased accuracy in subjects with the 9/10 genotype (i.e. high striatal dopamine levels; *M* = 60.00 ±5.36) compared to those with a 10/10 genotype (i.e. low striatal dopamine levels; 10/10 DAT1, *M* = 56.00 ± 6.51; *t*_(39.60)_ = 2.14, *95% CI* [0.21, 7.63], *p* = .04, *d* = 0.66, *BF*_*10*_ = 3.16), while this difference did not exist in the placebo group (9/10 genotype: *M* = 57.21 ±6.60; 10/10 genotype: *M* = 56.90 ±6.34; *t*_(30.01)_ = 0.15, *p* = .88). However, estradiol administration did not improve accuracy in subjects with the 9/10 genotype relative to placebo (9/10: *t*_(28.48)_ = 1.35, *p* = .19; 10/10: *t*_(40.11)_ = 1.79, *95% CI* [-0.41, 6.77], *p* = .08, *d* = 0.55, *BF*_*10*_ = 1.92).

We also predicted that the effect of estradiol would be moderated by subjects’ traits related to reward responsiveness such as the one measured by BIS/BAS ^30,31,33^. Indeed, when we accounted for subjects’ reward responsiveness (Fig 3C), we found that those who received estradiol (vs. placebo) were more accurate (β = 24.17 ±11.14, *F*_(1, 85)_ = 4.7, *p* = 0.03, Ω^2^ = 0.008), which was further moderated by reward responsiveness (*F*_(1, 85)_ = 4.6, *p* = 0.03, Ω^2^ = 0.036) while controlling for covariates. That is, estradiol enhanced accuracy specifically in subjects who were less reward responsive (*r* = −0.52, *p* < .001) with no such correlation in the placebo group (*r* = −0.16, *p* = 0.27). This result further supports the hypothesis that estradiol administration increased reward sensitivity by enhancing striatal reward prediction errors that increased the saliency of each trial ^8^. In our task, this hypothesis would predict more reward responsive subjects to switch between the choice options more frequently. Indeed, we found that within the estradiol group, there was a significant positive correlation between reward responsiveness and switching (*r* = 0.37, *p* = 0.009) that was only trending in subjects who received placebo (*r* = 0.26, *p* = 0.07). Importantly, the degree of switching negatively predicted subjects’ task accuracy (*F*_(1, 98)_ = 69.38, *p* < .001, Ω^2^ = 0.41) across both groups, explaining why more reward responsive subjects who received estradiol were less accurate.

### Increased reward sensitivity is observed in increased learning rates

Given our observation that estradiol increased reward sensitivity (Fig. 2), the interactive effect with DAT1 and reward responsiveness (Fig. 3), and our hypothesis that the differences in choice behaviour might occur due to upregulated striatal prediction errors, we predicted that estradiol would enhance the learning of reward probabilities. In a RL framework this would be reflected in increased learning rates.

To compare learning rates we used the maximum a posteriori estimates of the winning model fitted in a hierarchical Bayesian way ^48^, which included separate learning rates for positive and negative prediction errors, a temperature parameter, and an irreducible noise parameter (see Supplementary Materials for details on model selection and parameter recovery).

This model revealed that estradiol administration increased the learning rates for positive and negative prediction errors compared to placebo (α_Positive_: *M*_Estradiol_ = 0.47 ±0.22, *M*_Placebo_ = 0.37 ±0.24, *t*_(97.51)_ = 2.36, *95% CI* [0.017, 0.19], *p* = .02, *d* = 0.47, *BF*_*10*_ = 4.77; α_Negative_: *M*_Estradiol_ = 0.45 ±0.31, *M*_Placebo_ = = 0.27 ±0.25, *t*_(93.55)_ = 3.2, *95% CI* [0.068, 0.29], *p* = .002, *d* = 0.64, *BF*_*10*_ = 35.03, Fig. 4A). We also compared both parameters by computing 95% Highest Density Interval estimates ^49^ (Fig. 4D) with stronger evidence in favour of the negative learning rate 95% HDI [0.04, 0.39] being higher in subjects who received estradiol compared to the positive learning rate 95% HDI [-0.04, 0.32]. Contrary to our expectations, the observed main effect of estradiol was not moderated by the DAT1 polymorphism in case of either learning rate (α_Positive_: *F*_(1, 80)_ = 0.24, *p* = .89, α_Negative_: *F*_(1, 80)_ = 0.12, *p* = .73).

**Fig. 4.**
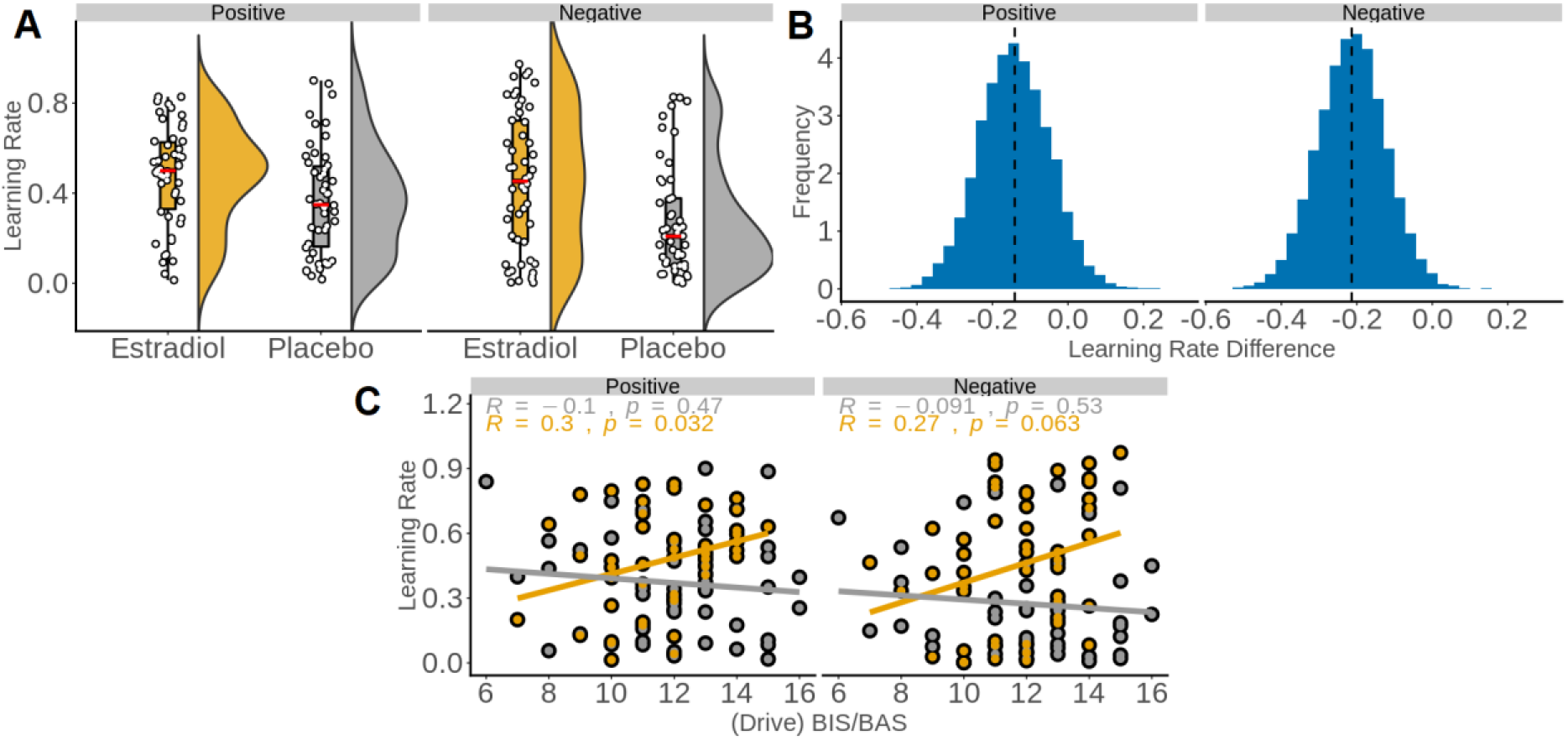
**A**) Learning rates by drug treatment. Individual dots represent subjects. The red bar represents the median, the box plot represents 75% of data, with the whiskers representing 1.95*IQR. **B)** 95% highest density interval (HDI) estimates for learning rate differences between both groups. **C)** Correlation between both learning rates and the drive subscale of BIS/BAS measuring motivation for goal-directed behaviour.

Finally, to provide construct validation for the obtained parameters, we correlated them with the BIS/BAS Drive subscale measuring motivation for goal-directed behaviour ^40^. We found that both positive and negative learning rates were weakly correlated in subjects who received estradiol (α_Positive_: r = 0.3, *p* = 0.03, α_Negative_: r = 27, p = 0.06) but not who received placebo (α_Positive_: r = −0.10, p = 0.47, α_Negative_: r = −.09, *p* = 0.53).

### Increased reward sensitivity is driven by differences in stay decisions, choice autocorrelation, and is moderated by the DAT1 genotype

Finally, to more precisely understand the observed difference in choice behaviour between both groups and DAT1 genotype, we tested whether these differences could be attributed to differences in staying behaviour and choice autocorrelation ^24,50^.

As would be predicted by increased accuracy in the 9/10 DAT genotype subjects, on average these subjects also chose the same option more often after being rewarded for this option before (*M* = 1.79 ± 0.18; Fig. 5A). This was true when comparing them to subjects with placebo who had the 9/10 genotype (*M* = 1.63 ± 0.22; *t*_(29.05)_ = 2.33, *95% CI* [0.02, 0.3], *p* = .03, *d* = 0.80, *BF*_*10*_ = 5.01), and when comparing them to subjects who had the 10/10 genotype (placebo: *t*_(41.61)_ = 2.22, *95% CI* [0.01, 0.27], *p* = .03, *d* = 0.66, *BF*_*10*_ = 3.34; estradiol: (*t*_(38‥86)_ = 2.49, *95% CI* [0.03, 0.27], *p* = .02, *d* = 0.64, *BF*_*10*_ = 5.95). In contrast, this difference was not observed between both drug groups when not accounting for differences in DAT1 genotype (*t*_(97.94)_ = 1.10, *p* = 0.28). In other words, the increase in accuracy by exogenously elevated estradiol in individuals with a 9/10 genotype was reflected in increased decisions to stay with options for which they were previously rewarded.

**Fig. 5.**
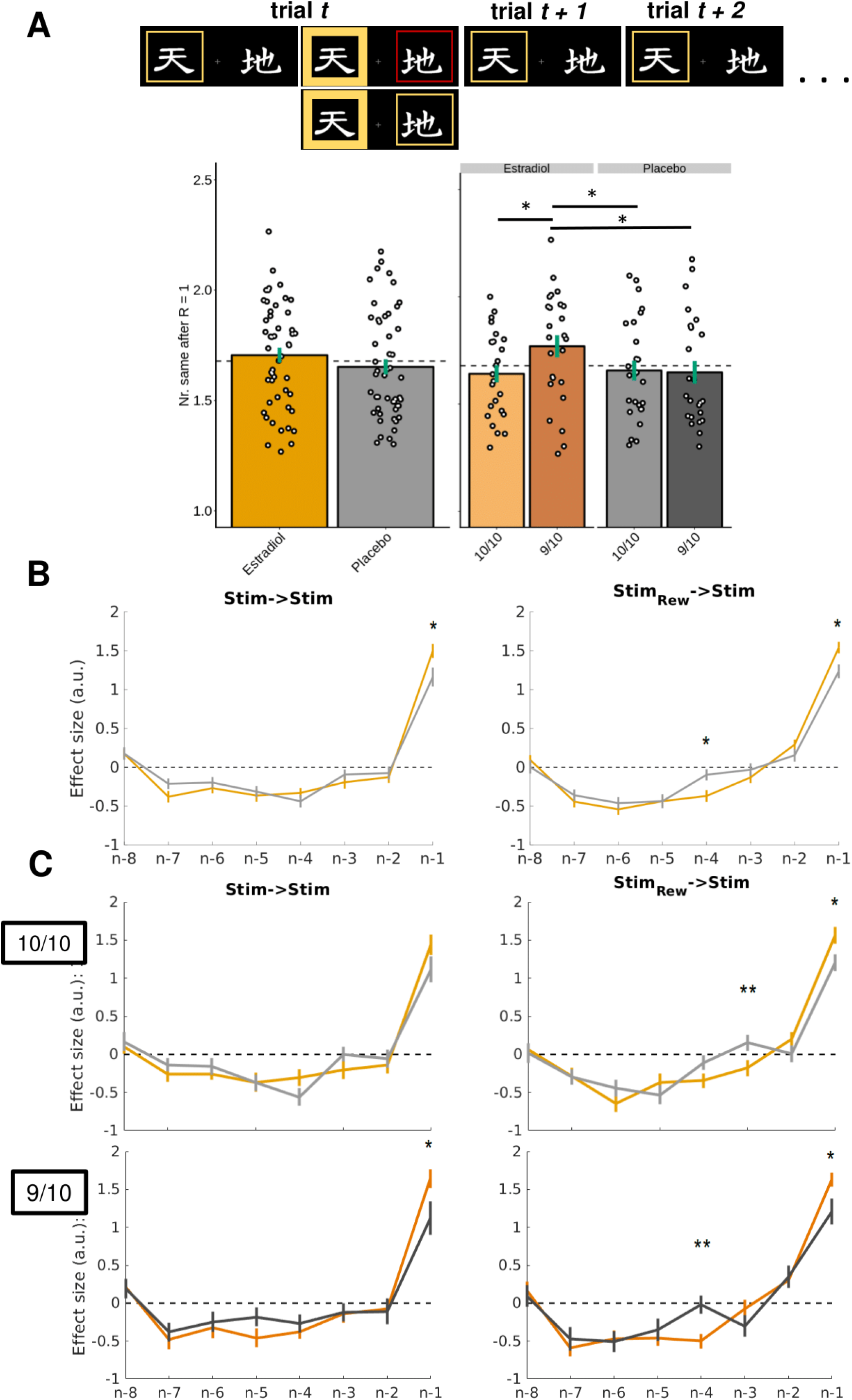
**A)** Staying behaviour: the average number of trials the same option was selected when that choice was previously rewarded. In both plots, each dot represents a subject, the green error bar represents standard error of the mean. **BC)** Autocorrelation analysis showing the impact of previous choices from 1 (n-1) to 8 (n-8) trials ago on the current choice. The left panel shows the averaged effect for both options irrespective of whether they were previously rewarded for that choice. The right panel shows the averaged effect for both options when they were previously rewarded for that choice. Both line plots are the mean and SEM of z-scored regressor weights. **B)** shows the split according to estradiol and placebo while **C)** shows a further split according to the 9/10 and 10/10 DAT1 polymorphism. ** *p* < .01, * *p* < .05.

We then extended this metric by examining how subjects’ previous choice impacted their current choice. We observed that estradiol administration caused subjects to weigh their choice on the previous trial (n - 1) more heavily compared to placebo (Fig. 5B), both when that choice was rewarded (*t*_(97)_ = 2.61, *95% CI* [0.073, 0.538], *p* = 0.01, *d* = 0.52, *BF*_*10*_ = 8.07) and irrespective of the outcome (*t*_*(97)*_ = 2.21, *95% CI* [0.034, 0.639], *p* = .03, *d* = 0.44, *BF*_*10*_ = 3.49). However, we also observed a decrease in the weight of previous rewarded choices occurring more than one trial ago (n − 4) (*t*_(97)_ = 2.59, *95% CI* [0.064, 0.482], *p* = 0.01, *d* = 0.52, *BF*_*10*_ = 7.93). In other words, while recent rewarded choices carried more weight, choices that were rewarded further in the past carried less weight, which is in line with having higher learning rates compared to the placebo group.

When we further split this analysis according to the DAT1 genotype (Fig. 5C), we observed similar results to the ones reported above. Estradiol administration enhanced the weight of the last choice (n - 1) on the current choice in the 9/10 subgroup (*t*_(32)_ = 2.12, *95% CI* [0.022, 1.027], *p* = .04, *d* = 0.73, *BF*_*10*_ = 3.44) while this was not the case in the 10/10 subgroup (*t*_(48)_ = 1.51, *p* = .14).

## Discussion

We examined the causal role of estradiol in reward processing in men. We show that estradiol affects reward processing and learning in men by increasing reward sensitivity, affecting choice autocorrelation, and increasing explicit awareness of changing reward probabilities. Furthermore, we show that estradiol’s effect on accuracy is moderated by subjects’ striatal dopaminergic functioning (DAT1 genotype) and reward responsiveness. Finally, we show that from a computational perspective, its effect can be characterised as an increase in learning rates.

First, we found that estradiol increased reward sensitivity during reinforcement learning. We observed that both groups systematically choose different options across trials, and on a subset of trials the options each group chose was statistically significantly different. Furthermore, estradiol increased choice autocorrelation for recent choices compared to placebo, while decreasing it for rewarded choice made several trials ago. These findings jointly show that by increasing reward sensitivity through upregulated striatal reward prediction errors^8^, estradiol administration caused recent choices and outcomes to be more salient compared to ones occurring further in the past. This interpretation is further supported by the estradiol group having higher learning rates for both positive and negative prediction errors, and subjects explicitly reporting to have observed a higher degree of reward probability changing. Furthermore, it is supported by the effect of estradiol on accuracy being moderated by reward responsiveness. The predicted increase in reward prediction errors caused more reward responsive subjects to switch more, in turn reducing their accuracy, while this relationship did not exist with placebo. The key contribution of the results above is showing, for the first time, the behavioural and computational effect of estradiol administration on reinforcement learning in men.

Next, we found that the effect of estradiol on accuracy was moderated by the DAT1 genotype, replicating previous correlational studies in women^6,75^. This result provides evidence for the hypothesis that estradiol may act by amplifying dopamine D1 receptor signalling in humans^2^, as such an effect would also have been predicted by dopamine precursor administration ^27 26,36^. To determine the robustness of this effect, we computed several behavioural metrics and Bayes factors to show converging evidence for a moderating role of DAT1. We found that estradiol administration significantly increased accuracy in subjects with the 9/10 (i.e. high striatal dopamine) compared to ones with the 10/10 genotype (i.e. low striatal dopamine) and that this increase in accuracy was likely driven by increased staying behaviour. That is, subjects with high striatal dopamine chose the same option on more trials, on average, if they were previously rewarded for that choice, compared to the other subgroups. Furthermore, even when they were not rewarded, they showed stronger choice autocorrelation compared to subjects with high striatal dopamine who received a placebo.

The key contribution of the DAT1-related findings is in reconciling discrepancies of previous correlational work that have been attributed to differences in baseline dopamine levels ^7,51–53^. That is, we show for the first time that baseline differences in dopamine, indeed, play an important role in how estradiol influences reward processing and that these need to be considered in future work aiming at better understanding their relationship. Crucially, the effects we report for accuracy and staying behaviour were not explained by other mechanistic explanations such as those related to androgen receptor functioning, androgen to estrogen conversion, or estrogen receptor functioning (see Supplementary Materials).

Our results highlight a further feature of how estradiol influences reinforcement learning. We found that estradiol improved accuracy specifically in the high striatal dopamine group which is in contrast to previous research with the dopamine precursor L-dopa where decreased accuracy in such subjects was reported but increased accuracy in subjects with low striatal dopamine was found ^26^. However, it is known that dopamine precursors impact behaviour in a dose-dependent manner ^54,55^. This would imply that our estradiol dose acted akin to a “low dosage” of a dopamine precursor which is supported by two estradiol administration studies in women. One showing 12 mg of estradiol (i.e. 6 times our dose) decreasing working memory performance^34^, which was interpreted as an overstimulation of dopaminergic transmission. With the other showing a dopamine-like dose-dependent effect of estradiol on hippocampal activity with doses between 2 and 12mg.^35^.

In addition to the effects reported for DAT1, we also provide preliminary evidence that the effect of estradiol is moderated by COMT (see Supplementary Materials). This is in support of the hypothesis that COMT activity is inhibited through estradiol metabolites which in turn increases dopamine availability ^25 2,56 13,57^.

The current study encountered limitations. We recommend future work employing pharmacogenetics to increase their sample size. While our sample size was approximately twice as large compared to most previous work ^7,25,34,36,52,53,58,59^ and in line with suggestions for the field ^2^, we suspect that an increase in sample size, as in work on other hormones ^60,61^, would have enabled us to make more precise claims at the level of polymorphism subgroup. As no previous estradiol administration study investigated either DAT1 or COMT, we had no basis for the minimal viable sample size, except the general recommendation in ^2^. Our results therefore await replication in future administration work.

### Conclusion

In conclusion, we have shown that estradiol influences choice behaviour by increasing reward sensitivity in healthy young men with distinct behavioural and computational signatures and that these effects are moderated by striatal dopamine-related genes (DAT1) and personality traits related to reward sensitivity. The approach and findings of this study show that understanding the role of estradiol in reward processing has important implications for a better understanding of the biology and neuroscience of human cognition that is moderated by genes in both health and disorder.

## Acknowledgements

The authors would like to thank Christina Faschinger and Isa Krol for their assistance in data collection, Nace Mikus for his help in data collection and analysis suggestions, and Lei Zhang for feedback on the manuscript. The study was supported by the Vienna Science and Technology Fund (WWTF VRG13-007).

## Supplementary Materials

## Methods

### Subjects

The short version of e-MINI ^62^ was used to screen and exclude those who had a non-diagnosed, disclosed, or a diagnosed psychiatric disorder. Subjects were recruited through social media, web portals, and flyers on university premises. All subjects provided written informed consent and were financially compensated for the completion of the experiment (50€) and received an additional maximum bonus of 40€ (range 7€ - 30€) based on their performance in the all the tasks.

### Measurement Instruments

#### Experimental Tasks

For each task, we gave subjects paper instructions including control questions to check whether all subjects understood the instructions. All tasks except for the N-BACK task were monetarily incentivised.

##### Reinforcement Learning

In addition to the main 500 trials, subjects were trained on 10 practice trials with two initial options to learn how the task works. The initial two options were then changed (i.e. the sensory cues) before the main trials started. We did this to avoid carry-over effects from practice to the main task. As shown in Fig. 1, each trial included three stages: (1) a cue onset stage (5 sec) where subjects had to decide between the two options and press the corresponding key. If they did not respond within that time frame, they would see a warning message indicating they should respond and try to be faster next time; (2) a choice feedback stage (1 sec) where subjects received information about both the chosen (thick frame) and unchosen (thin frame) option (yellow - correct, red - wrong); and (3) an inter-trial interval (*M* = 1.5 sec, jittered between 0.9 to 2.1 sec).

##### Working memory capacity

As described in the manuscript, subjects were presented with a sequence of letters one-by-one. For each letter, they had to decide if the current letter was the same as the one presented *N* trials ago by pressing “R”, in case it was not the same they had to press “O”. For example, in the 3-back condition, the letter sequence “***A B D A A***” would require subjects to press “R” only to the second occurrence of A, as this was the same letter as the one 3 trials ago. The last A in this example sequence is defined as a lure trial, while the other letters were nontarget trials. Lure trials were present only in the 2-BACK and 3-BACK conditions as in ^25^, and while lure trials were added to keep the task consistent with their implementation, we did not further analyse them separately as they were not relevant for our question. In total, there were four blocks per condition. Each block was announced by an instruction lasting for 2 sec (Fig. 1A), a fixation cross (1 sec) and a sequence of 20 trials. Each trial was presented for 1 sec with a 1 sec feedback phase and a 1 sec inter-stimulus interval. After every 20 trials, subjects had a 3 sec resting period, before the next block was announced. A lack of response to any cue was considered a miss.

### Procedure

During the first session, we assessed height, weight, abdominal, and visceral fat because these variables could impact estradiol metabolisation ^63,64^. On the second session subjects applied a topical transparent gel on their chest and shoulders that either contained 2 mg of estradiol (Divigel, Orion Pharma AG, Zug Switzerland) or a placebo. A male experimenter was present to ensure that the subjects applied the gel correctly.

### Statistical analysis

#### Behavioural analysis

To quantify statistical significance for the cumulative choice difference reported in the manuscript, we employed permutation testing (2000 iterations) by shuffling the responses of each subject and thereby decoupling the label from the responses, thus building a null distribution. The null distribution shows the difference that would be expected from random allocation to the group. To determine significance, we computed z-scores as measures of standardized effect size (e.g. ^65^) by subtracting from the quantity of interest the mean of the null distribution and dividing it by the standard deviation of the null distribution. From this, we were able to use the Fisher-z-transformation to obtain p-values.

We used two-sample proportion z-tests to determines the percentage of trials on which the number of subjects who chose option A in one group was statistically significantly different from the other group on a trial-by-trial basis.

For all general linear models reported in the main results, we regressed out several nuisance regressors known to impact estradiol metabolism or affect reinforcement learning behaviour. These included weight, BMI, abdominal and visceral fat ^63,64^, post-administration cortisol levels ^45^, and beliefs related to having received the drug. This was done because of our previous work showing the impact of beliefs about a hormone on subsequent behaviour, irrespective of whether subjects underwent treatment or received placebo ^66^.

All linear models were compared with BIC and AIC. Unless stated otherwise in the main text, for all reported results the winning model regressed out cortisol levels following administration, beliefs about having received the drug, the certainty of that belief and whether they had observed any changes in themselves, a composite score of weight and BMI that were summed together ^67^ because of their high intrinsic correlation (*r* = 0.89), visceral, and abdominal fat. For general linear models involving accuracy, we also regressed out reaction times to control for accuracy-speed trade-offs. All nuisance regressors were z-scored.

We compared both treatment groups for age and other bodily characteristics (i.e. BMI, height, weight, visceral, and abdominal fat) and potential differences in self-reported mood (MDBF), impulsiveness (BIS-11) and reward responsiveness (BIS/BAS) (see Questionnaires, Table S4 and S5). We used two-tailed independent samples Welch t-tests, or Wilcoxon signed-rank test if assumptions of normality were not met, to test whether the groups matched on all variables. To test for mood differences after administration between the treatment groups, we performed an ANCOVA for each of the three subscales of the MDBF questionnaire where we controlled for baseline mood scores. Two-way ANOVAs were further performed on the individual subscales of the BIS-11 questionnaire to investigate whether there was an interaction between the group (estradiol, placebo) and session (pre, post) on impulsiveness.

To compare working memory capacity assessed by the N-BACK task, we analysed target accuracy, reaction times, and d-prime. We analyzed this with an ANOVA containing the between-subject variable group (estradiol, placebo) and within-subject variable for condition together with an interaction term for group and condition.

In the supplementary results we used generalized linear mixed effects models using the lme4 package in R ^68,69^ to investigate whether the interaction between drug, DAT1 or COMT and trial would be predictive of the subjects’ chosen option.

#### Computational modelling

To test whether estradiol would increase reward sensitivity and thereby learning, we formalized behaviour within a reinforcement learning framework and fitted several Q-learning models ^70^ with softmax choice rules:

Q-learning model (equation 1):

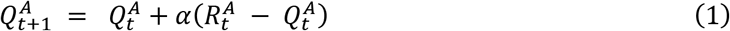

Softmax choice rule (equation 2):

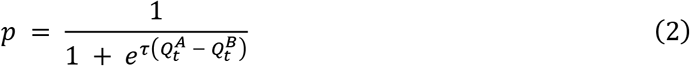

Where, *t* is time, *A* is option A, *Q* is subjective value, *α* is the learning rate, *R* is the obtained reward, and *τ* is the temperature parameter. Equations 1 and 2 represent our first model (model 1). In Q-learning, the basic idea is that agents learn subjective values of actions they perform in their environment. Subjective values are learned and updated through a value function (Equation 1) following feedback after each action. A teaching signal known as the learning rate-weighted prediction error dictates how strongly the subjective value will be updated on each action. The prediction error corresponds to the difference between the obtained and expected reward (i.e. the subjective value prior to making the new choice). Within this process, the learning rate dictates how heavily new information will be weighted in proportion to previous information about the option, and therefore how strongly the subjective value will change from its current estimate. The softmax equation then yields the probability of selecting an action given the learning rate and the temperature parameter, which reflects stochasticity of choice behaviour.

By employing computational modelling of this sort, we were able to obtain parameter estimates that quantify the difference in subjects’ behaviour, captured by a difference in learning rates. To obtain a more precise account of the effect of estradiol on reward processing, we extended the basic Q-learning model in several ways, as described below.

The first extension (equation 3a and 3b) allowed for separate learning rates for *α*_*Pos*_ and *α*_*Neg*_ that would differentiate between learning from positive and negative prediction errors.

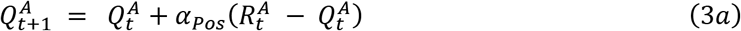

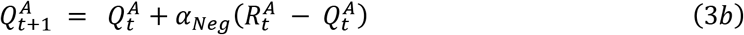

The updating with a positive learning rate occurs when the prediction error term 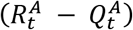 evaluates to positive while updating with the negative learning rate occurs when the prediction error term evaluates to negative. Furthermore, due to reward stochasticity of our task reward probability distribution (obtained by a Gaussian random walk - Fig. 1B), we added an additional parameter *ξ*, representing a lapse or irreducible noise parameter ^71^ in our choice rule (equation 4):

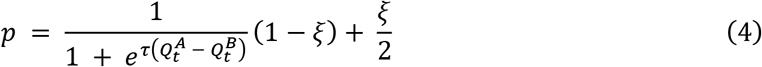

Finally, we added a perseverance parameter *λ* ^72^ (equation 5):

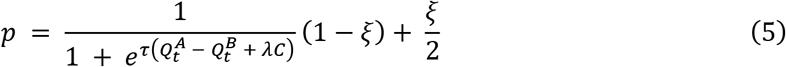

Where C = 1, if the same option was chosen on trial *n* and trial *n+1*, and C = −1 if the opposite was true. In summary, our full model had separate learning rates for positive and negative prediction errors, a choice stochasticity, irreducible noise, and perseverance parameter. All other models were reduced cases of this model and all possible combinations of the described free parameters therefore yielded eight models in total for which we estimated parameters:

**Table S0.** Models and parameters.

|  |  |
| --- | --- |
| Model 1 | Learning rate, temperature |
| Model 2 | Learning rate, temperature, lapse |
| Model 3 | Learning rate, temperature, perseverance |
| Model 4 | Learning rate, temperature, lapse, perseverance |
| Model 5 | Positive and negative learning rate, temperature |
| Model 6 | Positive and negative learning rate, temperature, lapse |
| Model 7 | Positive and negative learning rate, temperature, perseverance |
| Model 8 | Positive and negative learning rate, temperature, lapse, perseverance |

The model fitting was performed using JAGS and the rjags (v 4.9) package in R (v 3.6.0). Each model was run with 5000 samples each with 1000 burn-in samples on three chains. Priors over parameters and hyperparameters were set to default as described in ^48^. We computed the leave one out information criterion using the loo package ^73^ and used this metric to compare the models. Furthermore, we performed Bayesian model comparison by computing the (protected) exceedance probability ^74^ using the VBA toolbox ^75^ to determine the best model and compare its congruency with the LOOIC measure. Finally, we extracted the posterior predictive density for each subject as a measure of predictive power of the best model. This was then compared to the actual behaviour as a measure of static (accuracy collapsed across time) and dynamic (accuracy at each trial across subjects) predictive accuracy. From the obtained maximum a-posteriori estimates we then generated synthetic datasets for each subject and refit the model using synthetic data to assess parameter recovery. We did this by correlating our original and recovered parameters to determine whether the learning rate parameters of the winning model were correlated.

### Genotyping

#### DNA extraction and quantification

Buccal swabs were collected using sterile cotton swabs (Sarstedt AG, Germany). DNA was extracted from swabs using the QIAamp DNA Mini kit (Qiagen, Hilden, Germany) and eluted in a final volume of 50 μL of QIAamp buffer AE (Qiagen). Human nuclear DNA was quantified using the Applied Biosystems (AB) 7500 real-time PCR instrument (Thermo Fisher Scientific, Waltham, MA) and the Quantifiler Human Plus quantification Kit (AB) following manufacturer’s recommendations.

#### Typing of repeat length polymorphisms

Genomic DNA fragments that contain polymorphic repeat sequences were amplified in two separate reactions: i.e. a multiplex PCR (simultaneously targeting AR(CAG)n, DAT1 VNTR, Erα(TA)n and Erβ(CA)n) and a singleplex PCR (targeting solely AR(GGN)n), respectively.

The multiplex PCR was performed using 5 ng template DNA in a reaction mix (total volume of 25 μL) consisting of 1 × GeneAmp PCR buffer (AB), 0.25 mM each dNTP, 2.5 units AmpliTaq Gold polymerase (AB) and target specific primers (AR(CAG), DAT1, ERα and Erβ; including 5’-fluorescent-dye-labeled forward primers; details provided in Table 1). The following protocol was applied using the Veriti 96-well thermal cycler (AB): 35 cycles at 95 °C for 30 seconds, 55 °C for 1 minute, and 72 °C for 1 minute. Before the first cycle, an initial denaturation (95 °C for 5 minutes) was included, and the last cycle was followed by a final extension step at 72 °C for 45 minutes.

The singleplex PCR was conducted using 5 ng template DNA in a reaction mix (total volume of 20 μL) containing target specific primers (AR(GGN)n, details provided in Table S1)), 0.5 μL Phire Hot Start II DNA polymerase (Thermo Fisher) in 1 × Phire reaction buffer (Thermo Fisher). Amplification was carried out on the Veriti thermal cycler (AB) and included an initial denaturation step at 98 °C for 30 seconds, followed by 33 cycles of 10 seconds at 98 °C, 30 seconds at 60 °C and 30 seconds at 72 °C. The last cycle was followed by a final extension at 72 °C for 10 minutes.

Aliquots of PCR products were diluted with Hi-Di formamide (AB), mixed with internal lane standard LIZ 600 v.2 (AB) and separated on the ABI 3500 Genetic Analyzer applying standard conditions. The number of repeats predicted by the GeneMapper ID-X software (AB) was in full agreement to the actual repeats determined by direct sequencing of PCR products using the BigDye Terminator Sequencing Kit v3.1 (AB) in selected DNA samples.

**Table S1.**
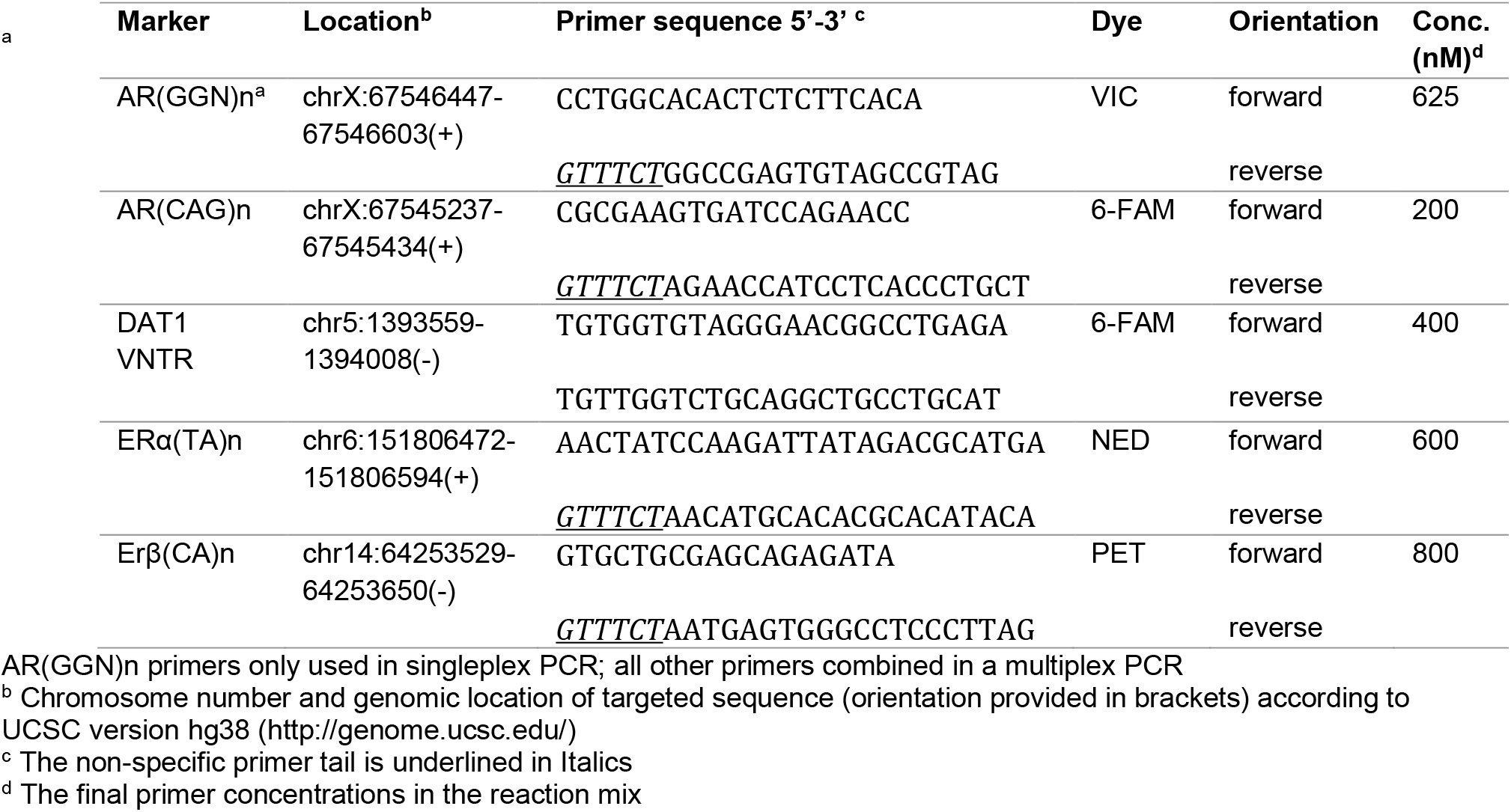
Panel of loci and primer sets used for the typing of repeat length polymorphisms.

#### Typing of the COMT Val158Met polymorphism

SNaPshot minisequencing was applied for the typing of Val158Met variants in the COMT gene. Therefore, a 177 bp fragment of genomic DNA harbouring the causative single nucleotide polymorphism (SNP rs4680) in its centre was amplified by PCR. The reaction mix comprised 5 ng template DNA, 1 × GeneAmp PCR buffer (AB), 0.25 mM each dNTP, 2.5 units AmpliTaq Gold polymerase (AB) and target specific primers (details provided in Table S2) in a total reaction volume of 25 μL. Thermal cycling was performed applying the Veriti cycler (AB) and conditions as follows: 95 °C for 5 min; 35 cycles of 95 °C for 15 seconds, 59 °C for 30 seconds and 72 °C for 1 minute; final extension at 72 °C for 5 minutes.

**Table S2.**
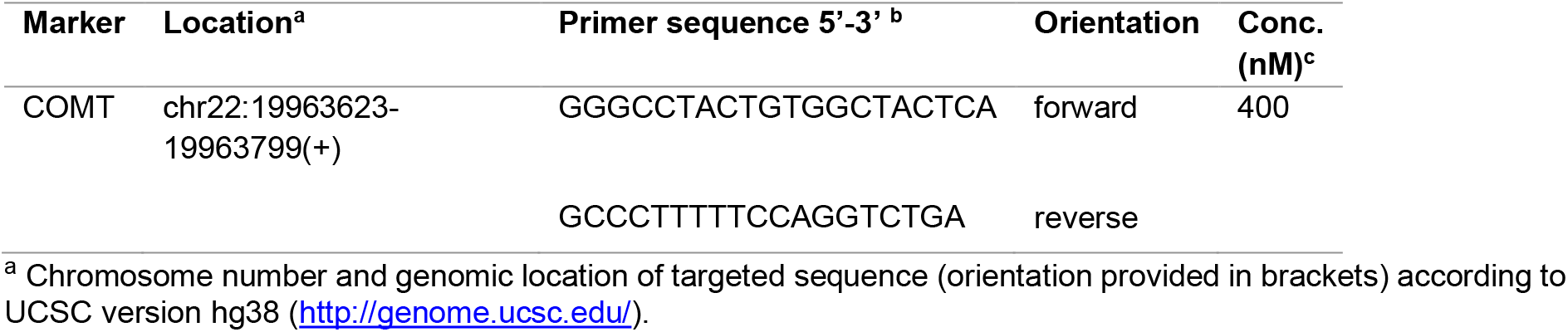
Primer set used for PCR of the COMT fragment.

PCR products were purified from excess primers and dNTPs by ExoSAP-IT (Thermo Fisher) treatment following manufacturer’s recommendations. Minisequencing was conducted on a Veriti thermal cycler (AB) in a total volume of 10 μL containing 3 μL of purified PCR product, 5 μL SNaPshot Multiplex Ready Reaction mix (Thermo Fisher) and 2 μL minisequencing primer (2 μM; details see Table 3). The cycling conditions (25 cycles) were as follows: denaturation at 96 °C for 10 seconds, annealing at 50 °C for 5 seconds and extension at 60 °C for 30 seconds.

**Table S3.**
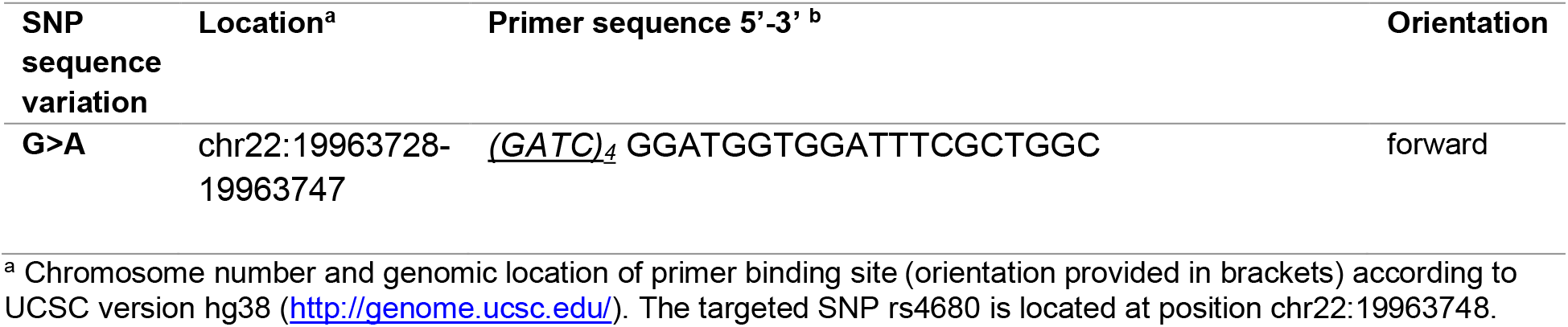
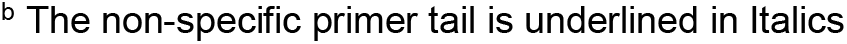
Minisequencing primer information.

ExoSAP-IT treatment was again applied for the clean-up of the minisequencing reaction. 5 μl of purified minisequencing reaction product was then mixed with 9.3 μL Hi-Di formamide (AB) and 0.2 μL of GeneScan-LIZ 120 internal size standard (AB). After a denaturing step for 5 min at 98 °C followed by cooling to 4 °C the fragments were separated on an ABI PRISM 310 Genetic Analyzer (AB) with POP4 polymer and analysed with GeneMapper v3.2 software. Calling of SNP variants based on minisequencing was in full agreement to results from direct sequencing of PCR products in selected DNA samples.

### Hormone concentrations

Quantification of estrone and estradiol in saliva samples was performed with derivatization using pentafluorobenzoyl chloride (PFBCl) and the addition of the isotopically labeled internal standards estrone-d_4_ and estradiol-d_5_. Organic saliva was reacted with 1.0 mL 1% PFBCl and 0.1 mL pyridine at 60°C for 30 min. The derivatization agents were evaporated, the sample was reconstituted with 0.5 mL NaHCO3 and extracted with 1 mL n-hexane. The organic phase was substituted with 0.2 mL dodecane and subjected to optimized GC-MS/MS analysis using an Agilent 7890 GC with Agilent DB-17ht 15 m × 0.25 mm × 0.15 μm capillary column connected to an Agilent 7010 tandem mass spectrometer operated in MRM mode using negative chemical ionization at 150°C with methane as a reaction gas (40%, 2 mL/min). Method validation was performed using ion transition m/z 464 -> 400 as a quantifier for estrone and m/z 660 -> 596 for estradiol, whereas a LLOQ of 1.92 fg o.c. and 1.94 fg was obtained, respectively.

Quantification of hydrocortisone and testosterone in saliva samples was performed using liquid chromatography tandem mass spectrometry (LCMS/MS), with an Agilent 6460 with electrospray ionization in positive mode coupled to a 1290 UHPLC system. Collision energy was optimized for specific MRM transitions of Hydrocortisone (363.2/121.1 m/z; 363.2/91.1 m/z), Testosterone (289.2/109.1; 289.2/97.1 m/z), 2,3,4-13C3-Hydrocortisone (366.2/124 m/z) and 2,3,4-13C3-Testosterone (292.2/100 m/z). Agilent Poroshell 120 EC-C18 was used for chromatographic separation under reversed phase conditions. The internal standard preparation and internal standard mixture was prepared containing 2,3,4-13C3-Hydrocortisone; 2,3,4-13C3-Testosterone, 2,4,16,16,17-d5-17b-Estradiol and concentration of 5ng/mL each. Samples were prepared by adding 100 μl internal standards (5 ng/mL) to 500μl plasma or saliva and the steroids were extracted using 4 mL MTBE. After 10 min. overhead shacking, the samples were centrifuged for 5 min. at 3000 rpm and the top MTBE layer was transferred to a test tube. MTBE was evaporated using a centrivap concentrator at 40°C (Labconco). The residual sample was then re-dissolved in methanol and analyzed by LC-MS/MS.

### Questionnaires

#### Mood

To control for a potential confound of mood, tiredness, or alertness from the treatment affecting subjects’ performance ^22^, we assessed subjects’ self-reported mood before and after administration of the treatment, using the German Multidimensial Mood State Questionnaire (“Der Mehrdimensionale Befindlichskeitfragebogen - MDBF) ^39^ *Both versions of this questionnaire (A and B) contain 12 items with a 5-level Likert scale and three subscales that test for different continuums of mood (Good-Bad [α*_*pre*_ *= .81, α*_*post*_ *= .77], Awake-Tired [α*_*pre*_ *= .84, α*_*post*_ *= .87], Calm-Nervous [α*_*pre*_ *= .73, α*_*post*_ *= .75])*.

#### Impulsiveness

We used the Barratt Impulsiveness Scale (BIS-11; ^40^ to measure subjects’ impulsiveness as ^52^ observed that variations in estradiol levels differentially affected women with low trait as opposed to high trait impulsiveness. BIS-11 is a widely used measure for impulsiveness with 30 items describing common behaviour and preferences related to (non)impulsiveness which individuals have to rate on a 4-point scale (1 - rarely/never, almost always/always - 4). The General Impulsiveness (α_pre_ = .71, α_post_ = .75) factor together with its three second-order factors (Motor Impulsiveness (α_pre_ = .47, α_post_ = .54) Nonplanning Impulsiveness (α_pre_ = .6, α_post_ = .63), Attentional Impulsiveness (α_pre_ = .49, α_post_ = .52) are reported.

#### Behavioural inhibition and activation

We measured the trait behavioural activation and inhibition with the Behavioural inhibiton/Behavioural Activation Scales (BIS/BAS; ^41^. The BAS scale is a 24-item questionnaire answered on a four-level scale (1- very true for me, 4 - very false for me). It is subdivided into Drive (α= .74), Fun Seeking (α= .67), and Reward Responsiveness (α= .6) while the BIS scale (α= .77) is unidimensional. Drive is thought to measure the persistent pursuit of goals (e.g. “I go out of my way to get the things I want”), Fun Seeking: the desire for new rewards and willingness to approach events that would be potentially rewarding (e.g. “I crave excitement and new sensations”), while Reward Responsiveness focuses on positive responses that would occur if a reward is anticipated (e.g. “When I am doing well at something I love to keep doing it”). Finally, the BIS scale measures sensitivity to negative events (e.g. “Criticism or scolding hurts me quite a bit”).

### Belief probes

In addition, we probed subjects’ beliefs and confidence about receiving estradiol (e.g. whether they believed they received estradiol or a placebo, how certain they were of this answer, and whether they noticed any changes). This was done to later regress out the potential contribution of beliefs arising, for example, from subjects researching potential side effects of the hormone prior the experiment. Namely, subjects’ beliefs about having received a hormone and beliefs about the effects of a hormone on their performance have previously shown to modulate behaviour independent of whether subjects had actually received it^66^.

## Results

### Matching of both groups

In the first part of the supplementary results, Table S4 and S5 show that our random assignment was successful as the groups did not differ in any of the measured parameters before (Table S4) administration and as a function of administration (Table S5). However, we did observe the expected change in estradiol metabolite concentrations in the estradiol group, outlined below.

### Hormone concentrations

We observed a statistically significant post-administration difference between both groups in log-transformed estradiol concentrations (*W* = 1545, 95% CI [0.03, 1.87], *p* < .05) with the estradiol group having higher estradiol metabolite concentration following administration (estradiol: *Mdn* = 41.77 ±531.54), placebo: *Mdn* = 5.55 ±230.23) but not before (estradiol: *Mdn* = 3.38 ±230.97), placebo: *Mdn* = 1.89 ±21.92) compared to the placebo group (*W* = 1498, 95% CI [-0.05, 1.03], *p* = .09). We report the median for the values above because even after log-transforming the metabolite concentrations, they were not distributed normally. Importantly, because we have observed high interindividual variance in estradiol concentrations prior to administration, we have reason to believe the obtained metabolite concentrations were contaminated during the handling of the samples following our data collection. Namely, in previous work such baseline variation was not observed despite an identical procedure and dosage with the main difference being that serum levels of estradiol were measured there ^38^. Log-transformed estrone and cortisol concentrations after administration were also examined showing no differences between both groups. Estrone: (experimental: *Mdn* = 8.79 ±4226.69), control: *Mdn* = 5.80 ±161.99) (*W* = 1427, 95% CI [-0.17, 1.05], *p* = .16), cortisol: (experimental: *Mdn* = 0.77 ± 0.94), control: *Mdn* = 0.73 ±1.15) (*W* = 1207, 95% CI [-0.31, 0.27], *p* = .90).

### Bodily measures and behavioural characteristics

As outlined in Table S4, both the estradiol and placebo group were also matched for their weight, height, BMI, visceral, abdominal fat, and individual sub scales of the BIS/BAS questionnaire (Drive, Reward, Fun-Seeking, Behavioural Inhibition). Similarly, separate one-way ANOVAs revealed no interaction for the four subscales of BIS-11 (Table S5) (General: *F*_(1, 195)_ = 0.01, *p* = 0.91, Attentional: *F*_(1, 195)_ = 0.04, *p* = .85, Motor: *F*_(1, 195)_ = 0.59, *p* = .45, nonplanning: *F*_(1, 195)_ = 0.08, *p* = .78).

Furthermore, we checked whether both the estradiol and placebo group did not differ in pre-existing differences in working memory (Figure S2A, S2B, S2C) in addition to testing whether administration influenced mood (Figure S2D). By doing so we were able to exclude differences in working memory and mood leading to the observed results ^26,76^. Separate ANCOVAs for the three subscales (Alertness, Mood, Calmness) of the MDBF revealed no differences in post-administration (Post) scores between the estradiol and placebo group when controlling for baseline scores (Pre) as a covariate (Mood: *F*_(1, 96)_ = 0.30, *p* = 0.58; Alertness: *F*_(1, 96)_ = 1.35, *p* = .25; Calmness: *F*_(1, 96)_ = 1.34, *p* = .25). Similarly, we observed no interaction between group membership and post-administration score (Mood: *F*_(1, 96)_ = 0.06, *p* = .81; Alertness: *F*_(1, 96)_ = 1.88, *p* = .17; Calmness: *F*_(1, 96)_ = 1.55, *p* = .22).

**Table S4:**
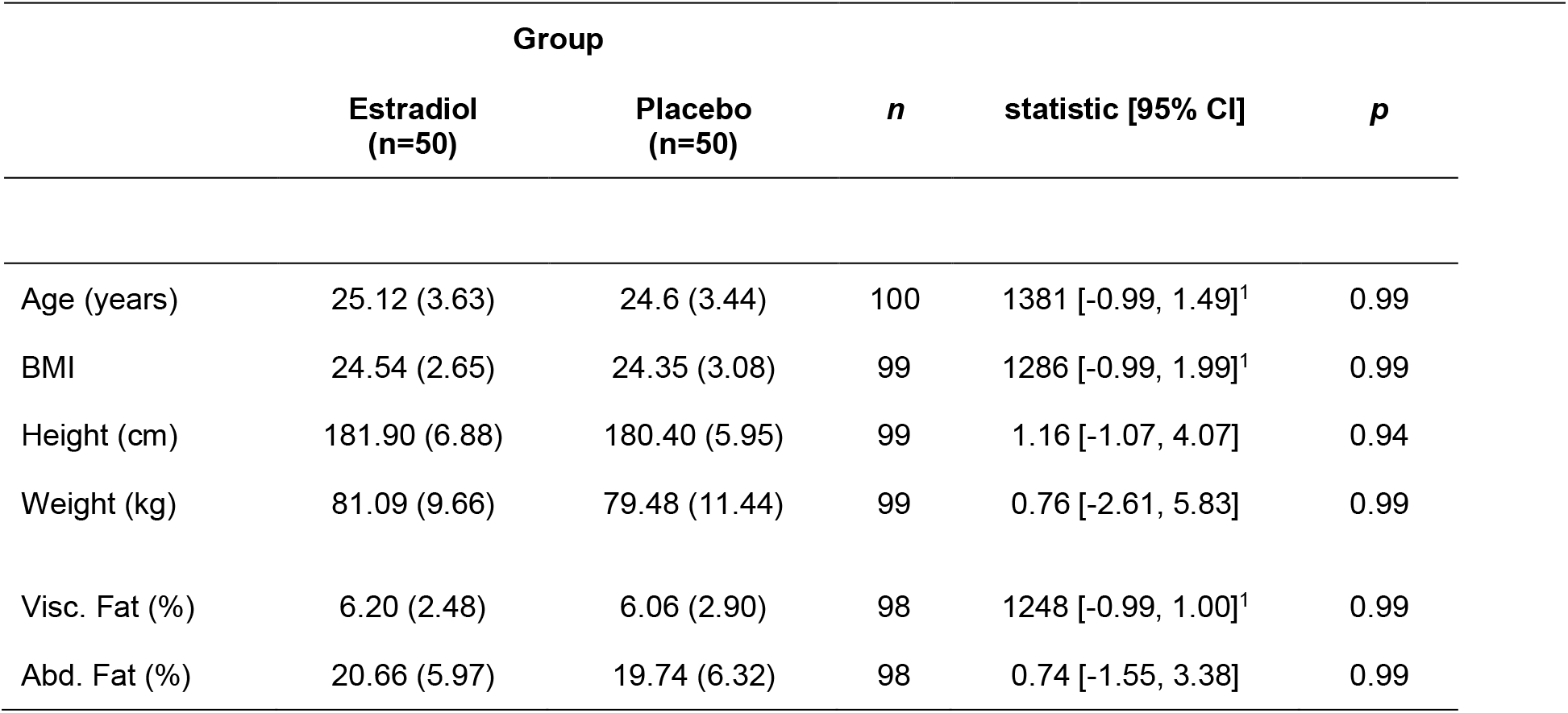
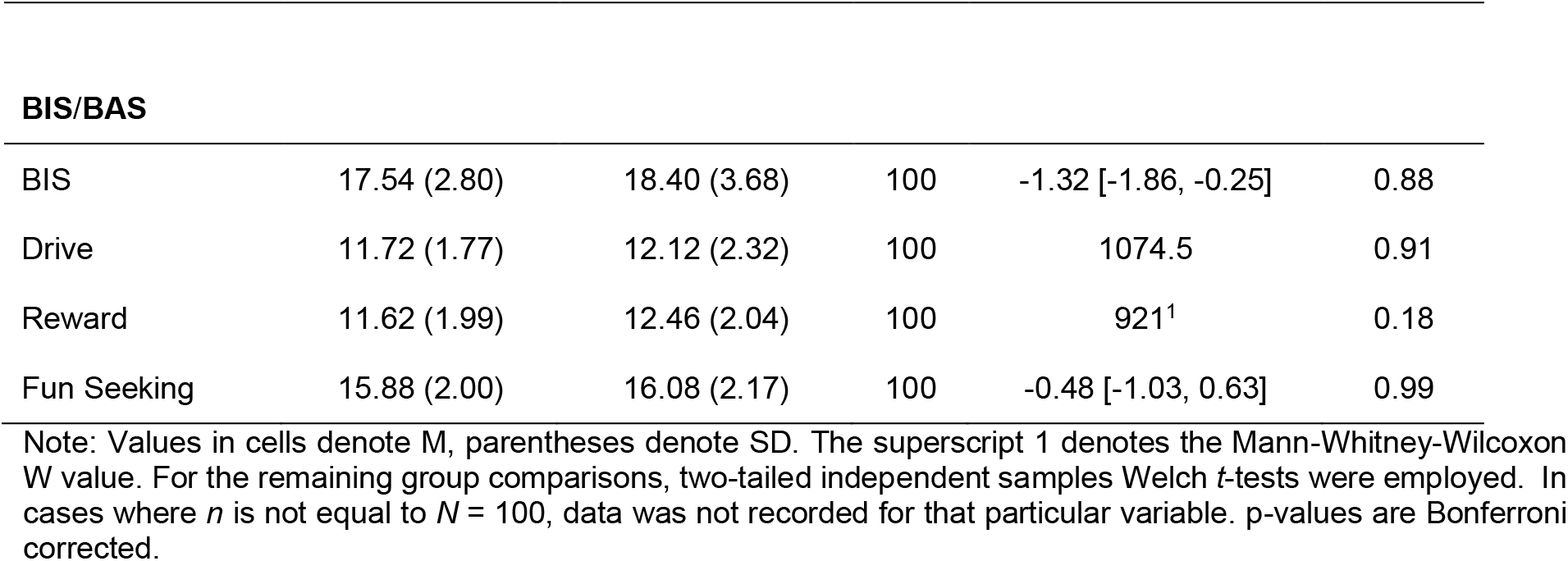
Descriptive statistics by treatment (Estradiol, Placebo).

|  | Group |  | *n* | statistic [95% CI] | *p* |
| --- | --- | --- | --- | --- | --- |
|  | Estradiol<br>(*n*=50) | Placebo<br>(*n*=50) |  |  |  |
| Age (years) | 25.12 (3.63) | 24.6 (3.44) | 100 | 1381 [-0.99, 1.49]<sup>1</sup> | 0.99 |
| BMI | 24.54 (2.65) | 24.35 (3.08) | 99 | 1286 [-0.99, 1.99]<sup>1</sup> | 0.99 |
| Height (cm) | 181.90 (6.88) | 180.40 (5.95) | 99 | 1.16 [-1.07, 4.07] | 0.94 |
| Weight (kg) | 81.09 (9.66) | 79.48 (11.44) | 99 | 0.76 [-2.61, 5.83] | 0.99 |
| Visc. Fat (%) | 6.20 (2.48) | 6.06 (2.90) | 98 | 1248 [-0.99, 1.00]<sup>1</sup> | 0.99 |
| Abd. Fat (%) | 20.66 (5.97) | 19.74 (6.32) | 98 | 0.74 [-1.55, 3.38] | 0.99 |
| **BIS/BAS** |  |  |  |  |  |
| BIS | 17.54 (2.80) | 18.40 (3.68) | 100 | -1.32 [-1.86, -0.25] | 0.88 |
| Drive | 11.72 (1.77) | 12.12 (2.32) | 100 | 1074.5 | 0.91 |
| Reward | 11.62 (1.99) | 12.46 (2.04) | 100 | 921<sup>1</sup> | 0.18 |
| Fun Seeking | 15.88 (2.00) | 16.08 (2.17) | 100 | -0.48 [-1.03, 0.63] | 0.99 |

Note: Values in cells denote M, parentheses denote SD. The superscript 1 denotes the Mann-Whitney-Wilcoxon W value. For the remaining group comparisons, two-tailed independent samples Welch *t*-tests were employed. In cases where *n* is not equal to *N* = 100, data was not recorded for that particular variable. p-values are Bonferroni corrected.

**Figure S1.**
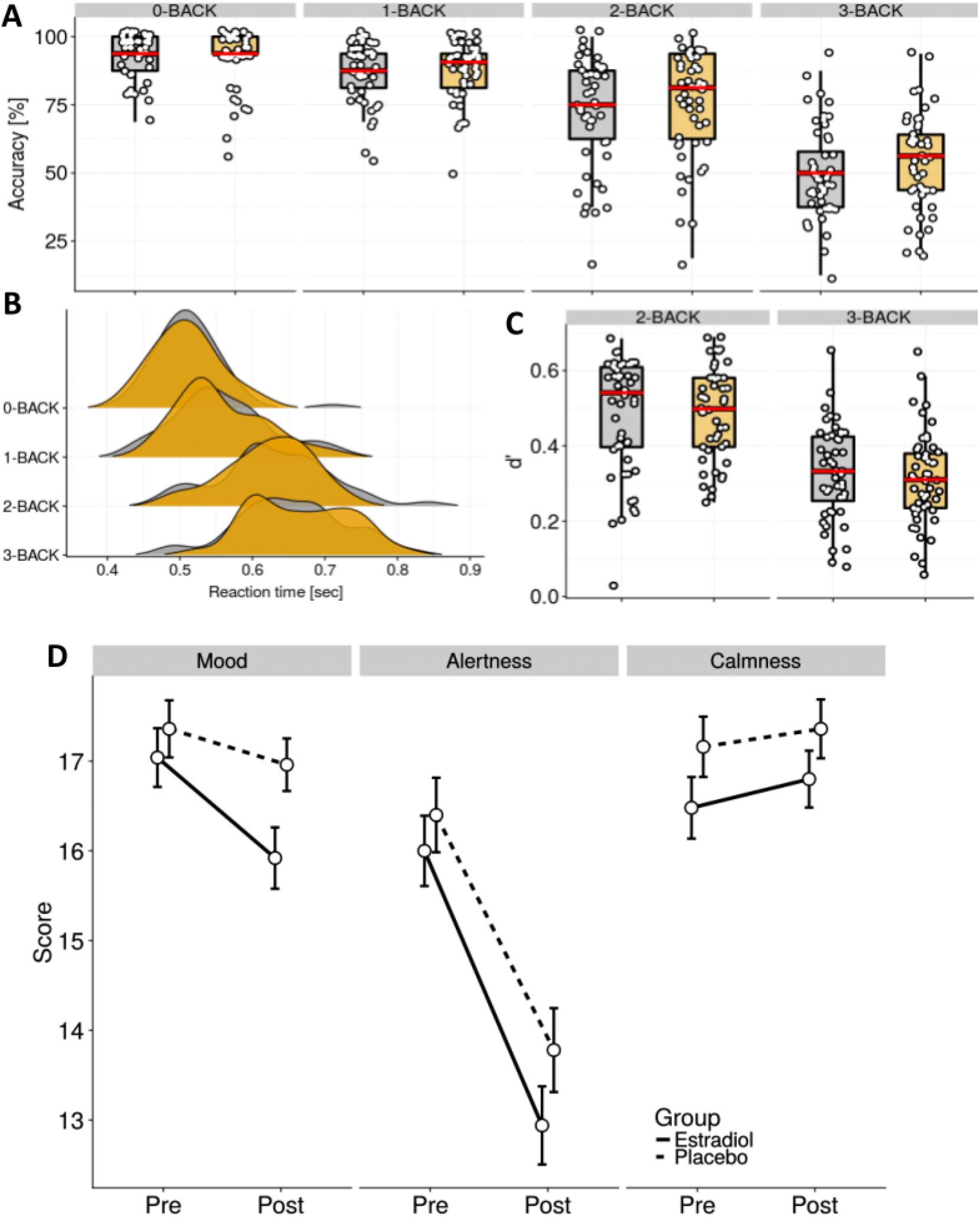
**A**) Accuracy for individual conditions. The red bar represents the median, the box plot represents the 75% middle most data points, with the whiskers representing 1.95*IQR. Orange depicts the estradiol and gray the placebo group. That is, they represent the division of subjects according to whether they would subsequently be allocated to the estradiol or placebo group. This color convention is used throughout all figures. **B**) shows density plots for reaction time data for individual conditions. **C**) shows d’ in the most difficult two conditions (2-BACK, 3-BACK) as there were no false alarms in the 0-BACK and 1-BACK, thus accuracy is reduced to d’. **D**) Average scores prior and post administration for the three subscales of the MDBF.

Furthermore, our working memory (N-BACK) task revealed a comparable picture for accuracy (Figure S2A), reaction times (Figure S2B), and d-prime (Figure S2C). That is, there was no statistically significant difference between the estradiol and placebo group in accuracy, average reaction times, and d-prime. We did observe an expected drop in performance in terms of decreased accuracy (0-BACK: 92.94 ± 9.34, 1-BACK: 88.06 ± 10.78, 2-BACK: 74.25 ± 19.38, 3-BACK: 51.56 ± 17.37), and d-prime (2-BACK: 0.48 ± 0.14, 3-BACK: 0.32 ± 0.12), and increased reaction times (0-BACK: 0.51 ± 0.05, 1-BACK: 0.56 ± 0.06, 2-BACK: 0.63 ± 0.07, 3-BACK: 0.66 ± 0.07) as the condition became more difficult (i.e. went from 0-BACK to 3-BACK). Separate linear models were used to compute to check for main effects of drug (*F*_(1,196)_ = 2.01, *p* = .16) and an interactive effect of drug and condition on d-prime (*F*_(1, 196)_ = 0.82, *p* = .37). As mentioned above, we also did this for accuracy (main effect of drug: *F*_(1, 392)_ = 1.07, *p* = .30; drug*condition interaction: *F*_(3, 392)_ = 2.30, *p* = .08), and reaction times (main effect: *F*_(1, 347)_ = 1.31, *p* = .25; drug*condition interaction: *F*_(1, 347)_ = 0.99, *p* = .39).

**Table S5.** Descriptive statistics of MDBF and BIS-11 subscales.

|  | *N* | Pre-administration |  | Post-administration |  |
| --- | --- | --- | --- | --- | --- |
|  |  | Estradiol | Placebo | Estradiol | Placebo |
| **MDBF** |  |  |  |  |  |
| Mood | 100 | 17.04 (2.31) | 17.36 (2.25) | 15.92 (2.41) | 16.96 (2.07) |
| Alertness | 100 | 16.00 (2.77) | 16.40 (2.93) | 12.94 (3.08) | 13.78 (3.30) |
| Calmness | 100 | 16.48 (2.43) | 17.16 (2.38) | 16.80 (2.24) | 17.36 (2.32) |
| **BIS-11** |  |  |  |  |  |
| General | 100 | 57.86 (7.68) | 59.28 (7.45) | 59.96 (8.51) | 60.46 (7.86) |
| Motor | 100 | 20.76 (3.00) | 21.78 (3.42) | 22.02 (3.42) | 22.42 (3.65) |
| Attention | 100 | 14.34 (3.37) | 14.14 (2.17) | 15.10 (3.30) | 14.64 (2.68) |
| Nonplanning | 100 | 22.76 (3.61) | 23.36 (4.29) | 22.84 (4.05) | 23.40 (4.21) |

Note: values in cells denote *M*, values in parentheses denote *SD*.

We observed no correlation between the certainty in the subjects’ belief as to whether they had received the drug or placebo (*r* = 0.02, *p* = .82), or between the reported observed changes and actually receiving estradiol (*r* = −0.08, *p* = .42). This shows that our double-blind procedure was successful, and that our placebo gel preparation was indistinguishable from the actual drug.

**Table S6.** Frequencies of individual polymorphisms of DAT and COMT genes.

| Polymorphism | Group | N |
| --- | --- | --- |
| 9/10 | Estradiol | 18 |
| 9/10 | Placebo | 16 |
| 10/10 | Estradiol | 21 |
| 10/10 | Placebo | 26 |
| Val/Val | Estradiol | 11 |
| Val/Val | Placebo | 9 |
| Met/Val | Estradiol | 23 |
| Met/Val | Placebo | 26 |
| Met/Met | Estradiol | 12 |
| Met/Met | Placebo | 15 |

Note: the split according to both COMT and DAT does not sum to 100 because for a few subjects it was not possible to determine their polymorphism.

### Reinforcement learning task

#### The role of CYP 19A1, ERα, ERβ, CAG, and GGN

Because the results for accuracy, staying, and switching behaviour that we report in the manuscript could also be moderated through other candidate mechanisms, we further analysed these candidate mechanisms together by providing theoretical motivation for the analyses. Here, we first briefly outline their importance and then summarize the observed results.

It is known that androgens are converted to estrogen ^77^. This means that the increase in estrogen levels arises from this conversion process and the administration more directly. Furthermore, variation in the length of two functional polymorphisms (CAG - polyglutamine, and GGN - polyglycine) are known to modulate the functioning of the androgen receptor gene ^78^. This is important for two reasons. First, our procedure has previously shown to increase circulating testosterone levels which could have raised estradiol levels whilst being moderated by subjects’ androgen receptor characteristics ^38^. Following from this, previous work has shown that brain regions important for memory and learning contain androgen receptors ^79^. Therefore, it could be possible that interindividual differences in both functional polymorphisms could have moderated our observed results due to interindividual variability. For example, greater CAG repeat length has previously been associated with lower scores in different cognitive tests in older men ^78^, and GGN repeat length has been associated with immediate and delayed logical memory recall in women ^80^. The described results show a correlation between individual variability in androgen receptor functioning and cognitive performance, giving rise to the possibility that CAG and GGN polymorphisms being potential candidate mechanisms moderating the observed effect of estradiol.

Throughout the conversion process from androgens to estrogens, the CYP19A1 gene encodes instructions for aromatase - the enzyme converting androgens to estrogens ^81^. The single nucleotide polymorphisms (SNPs) associated with the CYP19A1 gene regulate the metabolism of androgens and mediate brain estrogen activity. Two specific SNPs (rS700518, rs936306) have been previously shown to have a role in cognitive functioning in humans. For example, men with the homozygous AA allele have higher estradiol serum levels and greater bilateral posterior hippocampal gray matter volume compared to men with the homozygous GG allele ^82^. While other work has shown a differential impact of homozygous CC alleles versus homozygous TT alleles on episodic memory recall in women ^83^. Given that our procedure has previously shown to increase circulating testosterone levels and that polymorphisms of the CYP19A1 gene are known to have a role in cognitive functioning, we aimed to exclude the possibility that this may have driven our observed effects, and analysed both single nucleotide polymorphisms of the CYP19A1 gene.

Once androgens are converted to estrogens, estrogen action is mediated through the estrogen receptors (ERα, ERβ). Both receptors are widely distributed throughout the brain, including regions of importance for cognitive functioning and reward processing ^84^. So far, it has been shown that ERα is responsible for most of estrogen-related activation. For example, it has been shown that SNPs of ERα are related to Alzheimer’s disease and are associated with the likelihood of developing cognitive impairment ^85^. We have, therefore, focussed on two particular SNPs of ERα: rs9340799, rs2234693. In contrast, little is known of a potential impact of ERβ. As an exploratory measure, we have included repeats of this receptor in our analysis as well.

Of the described candidates (CAG, GGN, CYP 19A1, ERα, ERβ), no test revealed any effect of interest. There was no interaction between drug group (i.e. estradiol or placebo) and either the SNPs of ERα: rs9340799 (*F*_(2, 84)_ = 0.66, *p* = .52), rs2234693 (*F*_(2, 84)_ = 0.63, *p* = .53) in relation to accuracy. Furthermore, the same was true for the interaction between CAG repeats and drug group (*F*_(1, 87)_ = 0.45, *p* = .51), GGN repeats and drug group (*F*_(1, 87)_ = 1.31, *p* = .26), and SNPs of the CYP19A1 gene and drug group (rs700518 *F*_(2, 84)_ = 1.84, *p* = .15, rs936306 *F*_(2, 84)_ = 0.34, *p* = .72). In a final examination, we also looked at the repeats of ERβ to determine whether this could have driven any of the observed effects. However, this was not the case for either recorded variant of ERβ (ERβ1: *F*_(1, 87)_ = 0.02, *p* = .89, ERβ2: *F*_(1, 87)_ = 0.00, *p* = .96).

Identical results were obtained for switching behaviour. While we observed a statistically significant interaction between estradiol administration and the COMT polymorphism (see next section), this was not true for any of the other mechanistic explanations. That is, no model showed an interaction between drug group and either of the SNPs of ERα: rs9340799 (*F*_(2, 84)_ = 2.90, *p* = .06), rs2234693 (*F*_(2, 84)_ = 2.88, *p* = .06), CAG repeats (*F*_(1, 87)_ = 0.10, *p* = .76), GGN repeats *F*_(1, 87)_ = 1.32, *p* = .25), and SNPs of the CYP19A1 gene (rs700518 *F*_(2, 84)_ = 1.81, *p* = .17, rs936306 *F*_(2, 84)_ = 1.08, *p* = .35) in relation to switching behaviour. As in the case of accuracy, we also looked at the repeats of ERβ. Again, there was no statistically significant contribution to switching behaviour from this predictor for either recorded variant of ERβ (ERβ1: *F*_(1, 87)_ = 3.05, *p* = .08; ERβ2: *F*_(1, 87)_ = 0.96, *p* = .33).

We finally repeated the set of analyses for staying behaviour with no effects found of any of these variables: SNPs of ERα: rs9340799 (*F*_(2, 84)_ = 1.69, *p* = .19), rs2234693 (*F*_(2, 84)_ = 1.79, *p* = .17), CAG repeats (*F*_(1, 87)_ = 0.38, *p* = .54), GGN repeats *F*_(1, 87)_ = 0.30, *p* = .59), SNPs of the CYP19A1 gene (rs700518 *F*_(2, 84)_ = 1.27, *p* = .29, rs936306 *F*_(2, 84)_ = 0.59, *p* = .55), and variant of ERβ (ERβ1: *F*_(1, 87)_ = 1.35, *p* = .25; ERβ2: *F*_(1, 87)_ = 0.86, *p* = .36).

In brief, we have shown that the effects related to accuracy, switching, and staying reported in the manuscript did not depend on (1) the overall androgen receptor functioning assessed by CAG and GGN repeat polymorphisms, (2) interindividual variability in the androgen-to-estrogen conversion process via the CYP19A1 gene polymorphism, and (3) estrogen receptor polymorphisms (ERα) or repeats (ERβ).

### The effect of estradiol administration on choice behaviour is moderated by polymorphisms of both COMT and DAT1

We directly tested whether the effect of estradiol on choice behaviour is moderated by polymorphisms of dopamine-related genes (e.g. COMT, DAT1) by using generalized linear mixed models. We specifically tested whether the interaction between drug, polymorphism (COMT or DAT1), and trial are a significant predictor of the options they would choose.

Based on the inverted U-shape dopamine hypothesis ^46^, we predicted that estradiol administration would upregulate reward sensitivity in subjects with low prefrontal dopaminergic activity (i.e. Val/Val) but would not, or would even impair it, in those with high prefrontal dopaminergic activity (i.e. Met/Met). The model predictions support this hypothesis as it predicted that subjects with a Met/Val (*β* = 0.20 ± 0.04, *95% CI* [0.11, 0.28], *z* = 4.56, *p* < .001) and Val/Val genotype (*β* = 0.37 ± 0.06, *95% CI* [0.26, 0.48], *z* = 6.99, *p* < .001) were more likely to select option A as trials progressed when they received estradiol (Fig. S2, S4). Option A was the more rewarding option throughout the task (percent trials rewarded: *M*_optionA_ = 53.70%, *M*_optionB_ = 42.91%).

Based on the prediction that estradiol indirectly increases striatal dopamine levels, leading to higher reward prediction errors, we expected that subjects with the 9/10 genotype (i.e. high striatal dopamine) would select the higher value option more often, while this would be less often true for subjects with the 10/10 genotype (i.e. low striatal dopamine). This prediction was supported by the model showing that subjects with the 10/10 genotype with placebo (β = −0.12 ± 0.04, *95% CI* [−0.04, −0.20], *z* = −3.03, *p* < .01) were the most likely to select the lower valued option A throughout task progression, while estradiol administration dampened this slope in subjects with the same 10/10 genotype (Fig. S4). Results from both generalized linear mixed effects models showed that once individual variation was considered, the effect of estradiol administration on choice behaviour across trials was moderated by striatal (DAT1) and prefrontal (COMT) polymorphisms.

**Figure S2.**
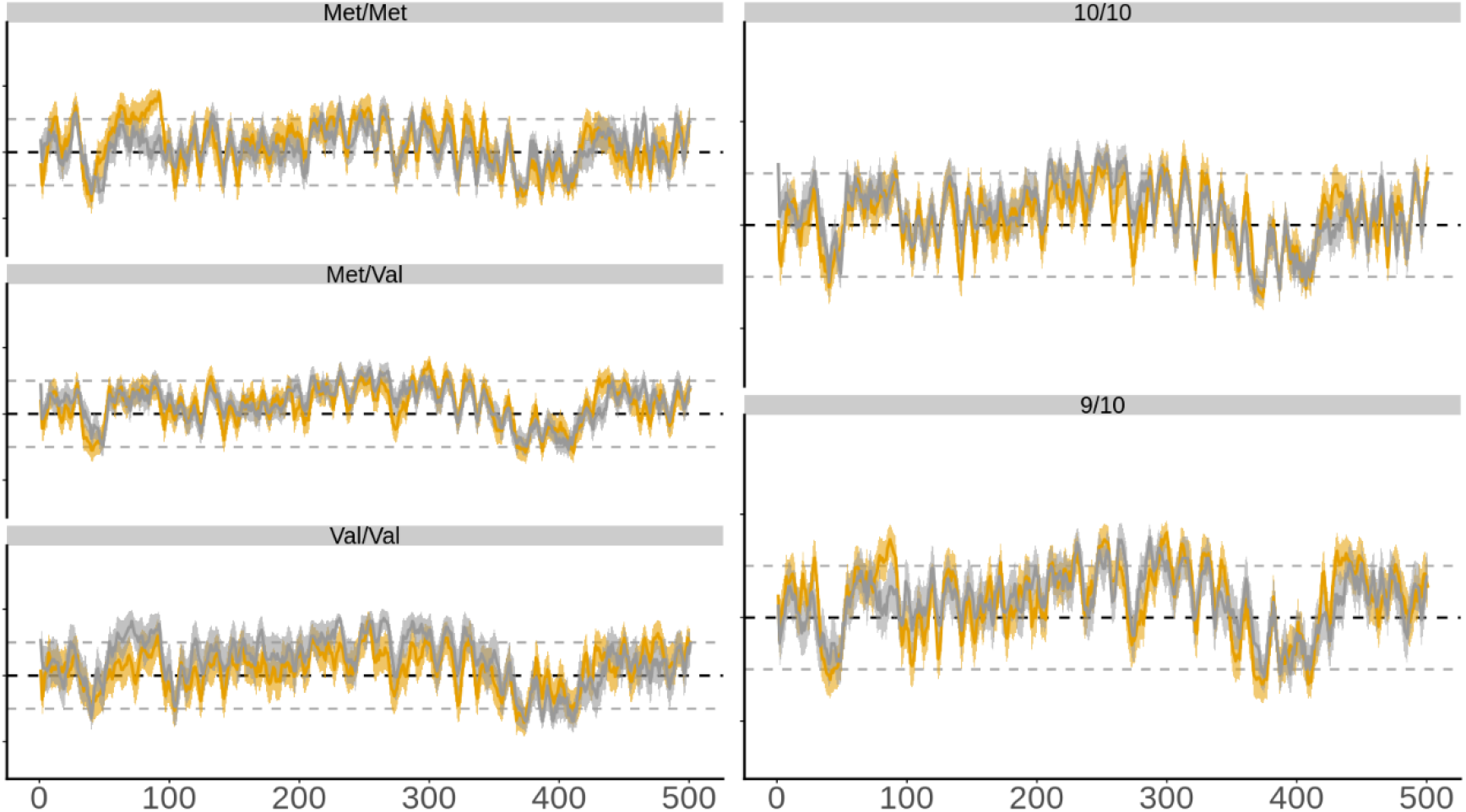
Relative choice probability for choosing option A (top of y-axis) vs. choosing stimulus 1 (bottom of y-axis) for the placebo (gray) and estradiol (orange) group split according to both polymorphisms assessed in the main text: COMT (left panel), DAT (right panel) across trials (1-500). Thick lines represent trial means, shaded areas denote standard error of the means. The blue line in the background denotes the empirical relative reward probability which was computed from the probability of stimulus two being rewarding (top of y-axis) - stimulus one being rewarding (bottom of y-axis). Gray dotted lines represent where subjects were on average 25% more likely to select option A (upper line) or stimulus 1 (lower line). All time-series traces are smoothed with a 5-trial moving average for visual purposes.

Figure S2 reveals a differential effect of estradiol administration on choice behaviour that depends on polymorphisms of both COMT and DAT. In the case of the COMT polymorphism this is most clearly visible in the lower left panel. The panel shows that placebo Val/Val subjects exhibited a clear tendency towards option A until trial ~370. After this, they did not reverse back towards choosing it more often despite option A being more rewarding from trial ~420 onwards. This is in contrast with results for subjects with other polymorphisms of COMT and results when subjects were split according to the DAT1 polymorphism. Estradiol Met/Met subjects exhibited choice behaviour more aligned with the reward probability distribution in the beginning at trial ~80 compared to subjects from the placebo group with the same polymorphism. When we then split subjects according DAT1 polymorphism, the estradiol 9/10 subjects can similarly be seen following the reward probability distribution more closely compared to the placebo 9/10.

Because of the results above and because estradiol administration likely results in increased prefrontal dopamine levels through downregulating COMT enzyme activity ^56^ we predicted that the interaction between estradiol administration and COMT polymorphism would be predictive of switching behaviour ^24^. As a measure of switching, we assessed the number of times the option chosen on trial *t* was different from the one chosen at trial *n + 1* (i.e. a switch), irrespective of the choice outcome on trial *t*. Estradiol administration did not significantly influence switch decisions (*M* = 162.12 ± 56.31) compared to placebo (*M* = 168.82 ± 68.13). However, we observed a significant interaction of estradiol administration by COMT genotype (*F*_(2, 80)_ = 3.22, *p* = .05, Ω^2^ = 0.04, Fig. S3). The interaction showed that subjects with placebo and a Val/Val genotype (i.e. low prefrontal dopamine availability) switched less often (*β* = −84.07 ± 33.69, p = .02) compared to all other groups. As predicted by the inverted U-shaped relationship between prefrontal dopamine levels and behaviour, Val/Val placebo subjects (Val/Val: *M* = 132.33 ±61.40) switched less compared to Met/Met placebo subjects (i.e. associated with high prefrontal dopamine availability; Met/Met: *M* = 204.27 ±53.52, *t*_(15.10)_ = 2.91, *95% CI* [19.25, 124.54], *p* = .01, *d* = 1.46). For the estradiol group, this difference was not present (Val/Val: *M* = 151.09 ±70.85; Met/Met: *M* = 178.5 ±55.34; *t*_(18.96)_ = 1.03, *95% CI* [-28.28, 83.10], *p* = .32, *d* = 0.44). In other words, estradiol administration attenuated naturally occurring differences in switching behaviour found in subjects with the Met/Met and Val/Val genotypes that are associated with high and low prefrontal dopamine levels, respectively.

**Figure S3.**
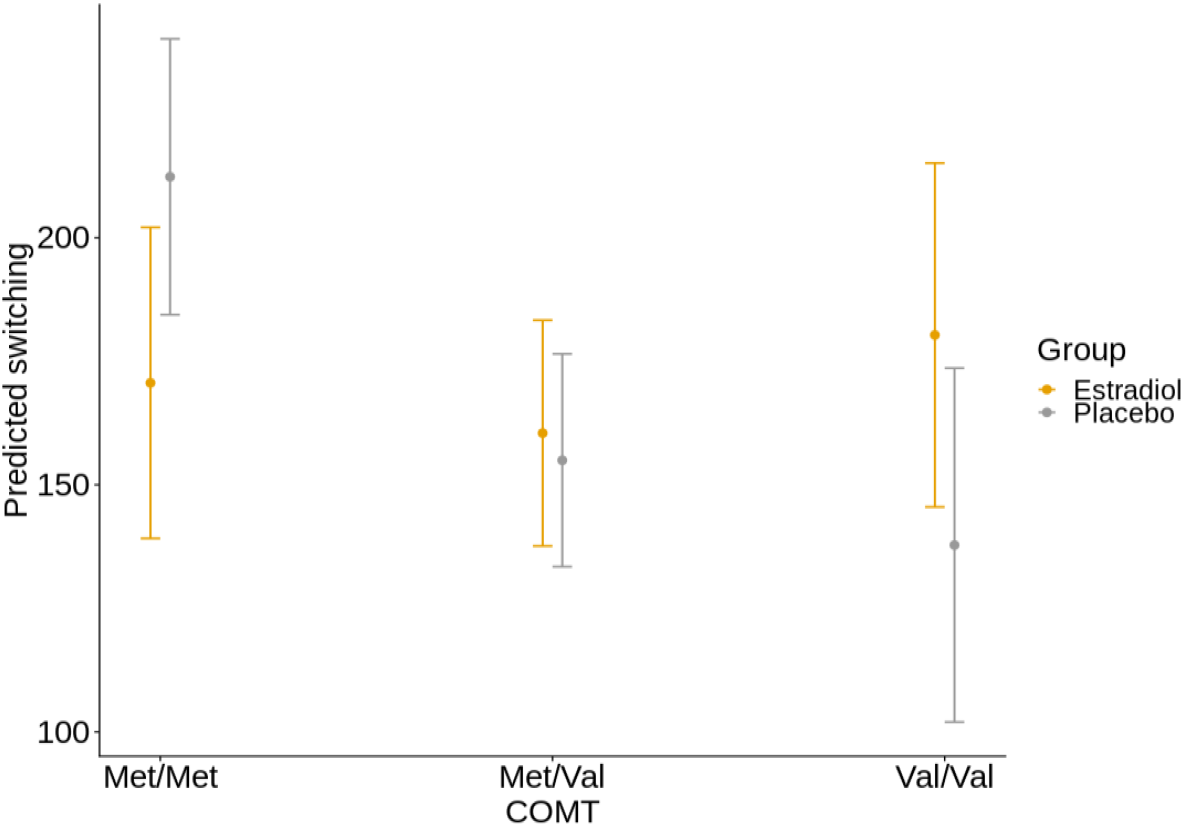
General linear model prediction for switching behaviour (i.e. a change in chosen stimulus on trial *t + 1* from trial *t*, independent of choice outcome on trial *t*). Estradiol administration dampened naturally occurring differences in switching behaviour when subjects were split according to the COMT polymorphism, i.e. whether subjects would switch the stimulus they chose on trial t compared to trial *t + 1* irrespective of choice outcome on trial *t*. Error bars represent SEM.

**Figure S4.**
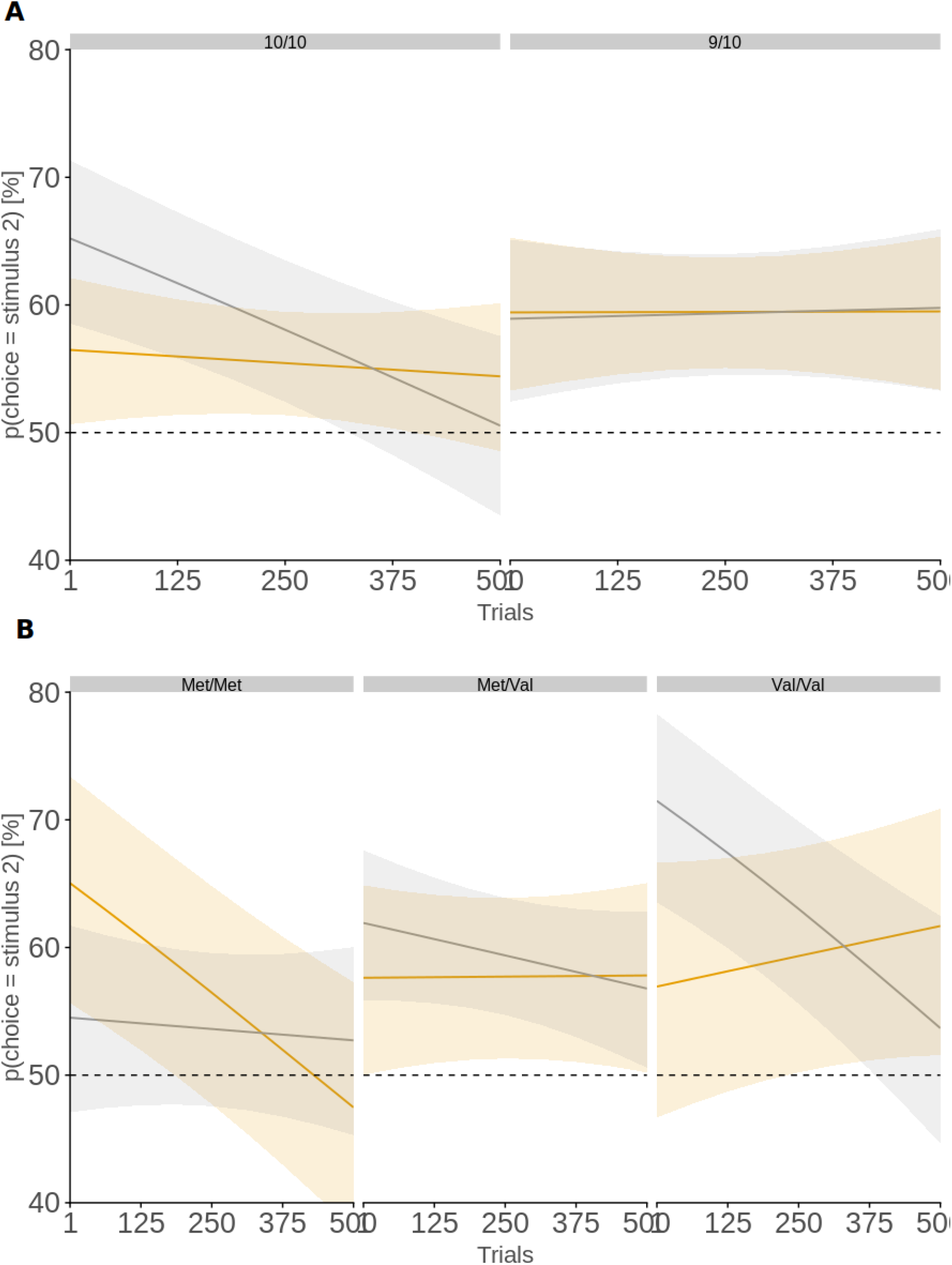
Predictions from winning models of the generalized linear mixed effects models for **A**) the interaction between drug, DAT, and trial on choice, and **B**) the interaction between drug, COMT, and trial on choice.

### Formal model comparison

We estimated parameters by fitting several Q-learning models. The best model (model 6, leave one out information criterion (LOOIC) = 58888, Fig. S5A) included separate learning rates for positive and negative prediction errors, a temperature parameter, and an irreducible noise parameter. The model predicted choice behaviour above chance (*t*_(99)_ = 13.95, 9*5% CI* [0.64, 0.68], *p* < .001, Fig. S5B) and performed equally well for both groups (*M*_Estradiol_ = 66.26 % ±10.77, *M*_Placebo_ = 64.90 % ±11.85; *t*_(97.115)_ = 0.76, *95% CI* [-0.03, 0.06], *p* = .45).

**Figure S5.**
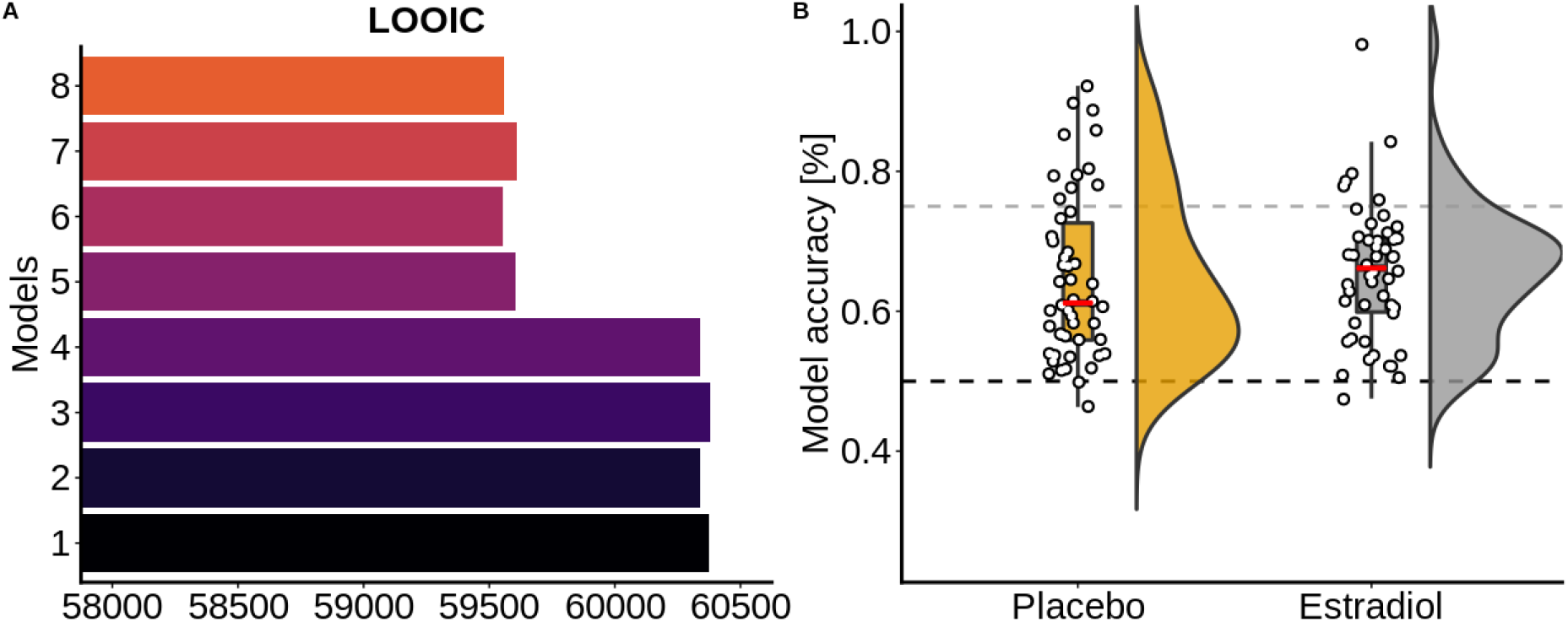
**A**) Leave one out information criterion (LOOIC) value for all employed models. Lower LOOIC indicates better model fit - model two was selected as the best model. **B)** The overall model accuracy collapsed over time obtained from the posterior predictive density shown for both drug groups separately.

In addition to computing the leave-one-out information criterion to perform model comparison ^73^ we similarly computed the exceedance probability of the winning model using the VBA toolbox ^75^. This value showed a strong preference for the winning model *P*(model two) = 98%. Furthermore, we computed protected exceedance probability ^74^ as an extension which, while yielding an expected decrease in the winning model probability, still favoured model two over other competing models (*P*(model two) = 12.5%). The likely decrease was due to the reinforcement learning task not being optimized to detect behavioural differences between the models tested. However, in all reported models, the latent variable of interest, i.e. the learning rate, remained unaltered. We would therefore expect the increase in learning rates to be present if we were to select the learning rates from models that best fit individual subjects.

### Validating model

We further tested the model validity and predictions by computing posterior predictive densities, i.e. what predictions does the model make on a trial-by-trial basis for subjects with the parameters such as those that were extracted from our subjects. Posterior predictive densities showed no difference in a fit between both the estradiol and placebo group and approximated the empirical reward probability distribution (Fig. S6A). To quantify this, we then compared model predictions from posterior predictive densities with actual subject behaviour to assess model accuracy collapsed across time (Fig. S6B) showing it performed above chance and equally well for both groups. We further compared accuracy on each trial across subjects to ensure that there were no unexpected drops in accuracy. This did not happen as the model (Fig. S6B) had no discernible drops in performance. We also performed parameter recovery on the winning model where we used the maximum a posteriori estimates reported in the manuscript, generated data for synthetic subjects, performed parameter estimation on the synthetic data, and correlated the newly obtained synthetic parameters with original parameters for each subject. This procedure showed that both original and recovered learning rates correlated with one another (negative: *r* = 0.33, *p* < .001, positive: r = −0.34, p < .001).

**Figure S6.**
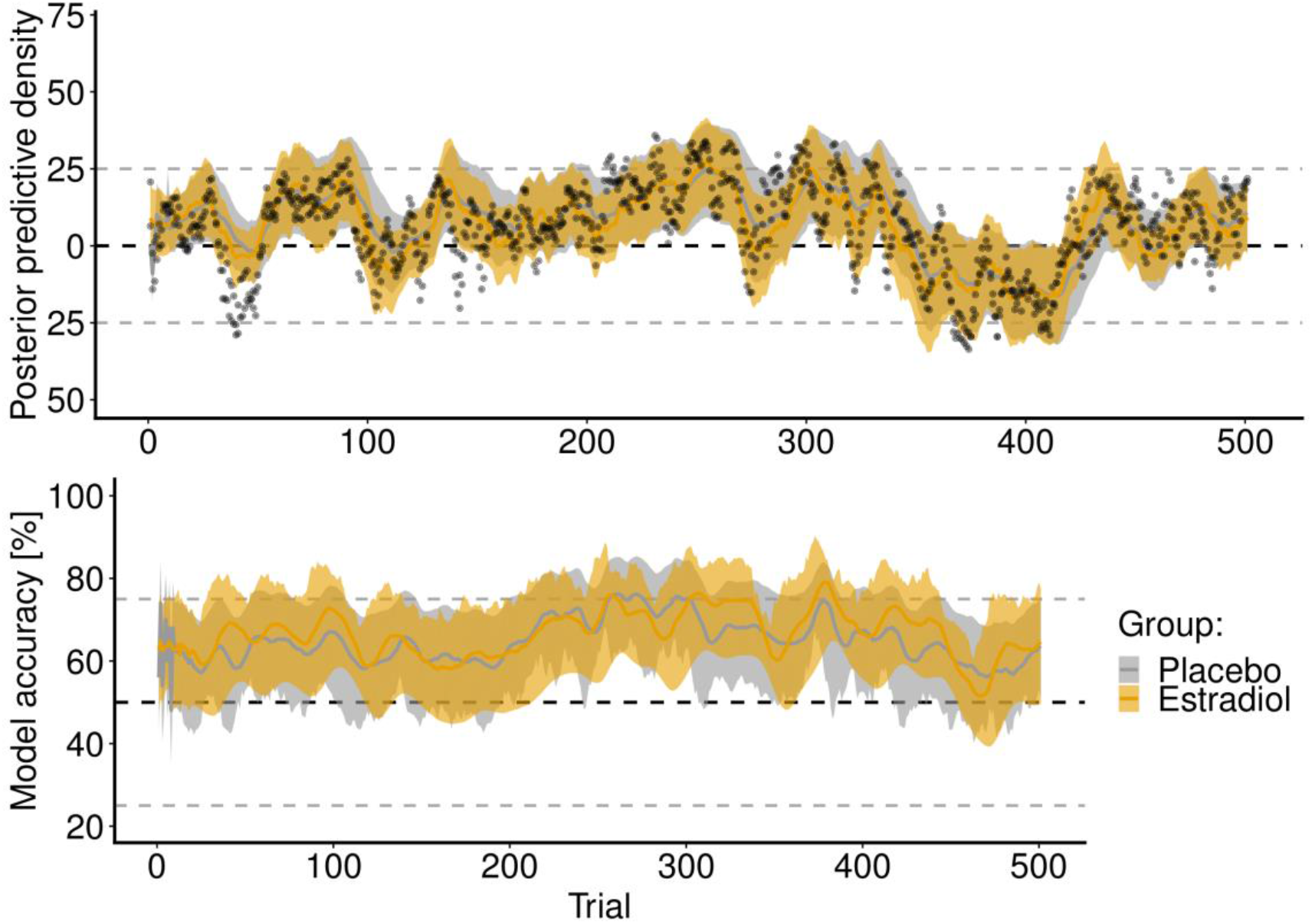
**A**) Posterior predictive density computed for both drug groups with overlaid average responses for both drug groups across trials **B**) Accuracy for both drug groups obtained from the posterior predictive density for both drug groups separately.

